# Motor circuit function is stabilized during postembryonic growth by anterograde trans-synaptic Jelly Belly - Anaplastic Lymphoma Kinase signaling

**DOI:** 10.1101/841106

**Authors:** Phil-Alan Gärtig, Aaron Ostrovsky, Linda Manhart, Carlo N. G. Giachello, Tatjana Kovacevic, Heidi Lustig, Barbara Chwalla, Sebastian Cachero, Richard A. Baines, Matthias Landgraf, Jan Felix Evers

## Abstract

The brain adapts to a changing environment or growing body size by structural growth and synaptic plasticity. Mechanisms studied to date promote synaptic growth between partner neurons, while negative counterparts that inhibit such interactions have so far remained elusive. Here, we investigate the role of Jeb-Alk signaling in coordinating motor circuit growth during larval stages of *Drosophila*. We quantify neuronal growth dynamics by intra-vital imaging, and synaptogenesis at nanometer resolution using endogenously labeled synaptic proteins, conditionally tagged with a fluorophore, and link changes in circuit anatomy with altered synaptic physiology and behavior. We find that loss of Jeb-Alk signaling leads to increased strengthening of synaptic excitation by developmental addition of additional postsynaptic but not pre-synaptic specializations. These changes ultimately lead to an epilepsy-like seizure behavior. We thus demonstrate that trans-synaptic anterograde Jeb-Alk signaling acts to stabilize developmental plasticity and circuit function, and that it does so specifically during postembryonic growth.

## Introduction

Neuronal circuits remain plastic and are continuously modified after birth in response to environmental change, thus supporting learning and memory, adaptation to a growing body or compensation for injury. To identify the molecular mechanisms that maintain nervous system function during adult life, but at the same time allowing neuronal circuits to stay plastic is, therefore, important.

During phases of initial circuit formation and subsequent circuit plasticity, axonal and dendritic terminals are decorated with thin cellular protrusions, termed filopodia ^1–3^. Filopodia undergo extensive dynamic outgrowth and retraction, effectively increasing the chance of pre- and postsynaptic arbors interacting, and thus of synaptogenesis. Observations in dissociated neuron culture suggest that dendritic filopodia, upon axonal contact, induce the formation of presynaptic specializations ^4^; and equally within axonal filopodia, presynaptic release sites differentiate following contact with postsynaptic targets ^5–8^. Such interactions between synaptic partners include non-neuronal cells, for example at neuromuscular junctions: filopodia from muscles (myopodia) are thought to aid partner selection through interactions with motorneuron growth cones ^9^. Emerging immature circuits are adjusted through activity-dependent mechanisms, leading to selective stabilization of appropriate neuronal branches and synapses, while other synapses are dismantled and their branches retracted ^10–13^. This initial phase of highly-plastic circuit wiring ultimately generates patterns of synaptic connections that support future network function, and which are subsequently maintained, even thrugh phases of considerable growth. For example in *Drosophila*, synaptic patterns between identified partner neurons that were established during embryogenesis are maintained as the animal grows, scaling several-fold in synapse number, proportionally with neuronal arbor size ^14–16^. The trans-synaptic signaling mechanisms that coordinate such scaling of pre- and postsynaptic partners during postembryonic circuit expansion are not known.

To date, pathways regulating the development of synaptic connectivity have primarily been studied during early stages of circuit formation, when synapses are first established and adjusted. Much less is known about later circuit growth. Here we focus on postembryonic stages in the *Drosophila* larva, during which neuronal growth and connectivity scale with substantive increases in body size, as also seen in vertebrates ^14–18^. Specifically, we identified Jelly Belly (Jeb) – Anaplastic Lymphoma Kinase (Alk) signaling, as a mechanism to stabilize growth of neuronal networks. Previously shown to regulate circuit formation in *Drosophila*, Jeb is released from axonal terminals, and detected by the receptor tyrosine kinase, Alk ^19–21^. In the visual system of the adult fly, Jeb-Alk signaling controls axonal morphology ^20^ and neuronal survival ^21^. At the larval neuromuscular junction this pathway negatively regulates synaptic coupling strength ^19^. On the functional level, loss of Alk signaling can express as increased memory performance in adult flies ^22^, mediated by dis-inhibition of protein synthesis ^23^. However, the cell-biological and structural processes that underlie these alterations in network performance are not known.

Using novel techniques to label endogenous proteins, or conditionally remove gene expression by mutation in individual cells, we confirm that Jeb is a presynaptic anterograde signal that is detected by Alk on postsynaptic dendrites in the motor system. Intra-vital imaging of neuronal growth dynamics, and quantification of endogenous synaptic protein accumulation reveal that Alk activation in the postsynaptic compartment inhibits the developmental proliferation of postsynaptic, but not of presynaptic specializations independent of dendritic arbor size. Alk activation in dendrites also negatively regulates presynaptic filopodial dynamics via retrograde feedback. Performing intracellular patch clamp electrophysiology, we find that Jeb-Alk signaling deficient nervous systems undergo increased developmental strengthening of synaptic coupling, ultimately destabilizing circuit function as evident by epilepsy-like seizures evoked by electroshock.

Our data show that during postembryonic growth of neuronal circuits, presynaptic filopodia enhance the growth of prospective postsynaptic partner dendrites towards presynaptic sites. Once contact is established, trans-synaptic Jeb-Alk signaling limits further circuit growth and, thus, is essential to stabilize circuit function.

## Results

### Subcellular localization of Jeb and Alk in the fly motor system suggests a synaptic anterograde signaling pathway

The regulation of synaptic coupling strength is essential for nervous system formation and function. At the larval NMJ, Jeb was previously identified as instrumental in setting synaptic coupling strength^19^. In adult flies, the Jeb receptor Alk had been shown to impinge on memory performance ^22, 23^, suggesting potentially a similar role for Jeb-Alk in the CNS. To investigate this, we generated novel genetic reagents with which to visualize the subcellular localization of endogenous Jeb and Alk proteins; and for conditionally removing gene function in single identified cells within the nervous system.

Jeb and Alk are prominently expressed in the larval CNS ^24–26^ (Fig S1). To identify the cell types expressing Jeb, we used a trojan GAL4 inserted into the Jeb locus (*jeb-T2A-GAL4*) ^27^ to report Jeb transcriptional activity, e.g., via a *UAS-nlsGFP* reporter. We find nlsGFP expression in neuronal cell bodies, but not glia cells (Fig S1B-G). To reveal subcellular Jeb distribution, we performed anti-Jeb immunohistochemistry on Jeb null mutant embryos in which we reinstated Jeb expression in a specific pre-motor interneuron IN_lat_, and two sensory neurons, ddaD and ddaE (genotype: *jeb^2^/Df(jeb); eyg-Gal4, UAS-jeb*) (Fig 1A-B). This revealed that Jeb accumulates at presynaptic release sites of both IN_lat_ and sensory neurons, as evidenced by co-expression of the active zone marker Brp^RFP^ (Fig 1B), though not in dendrites (data not shown).

**Figure1.**
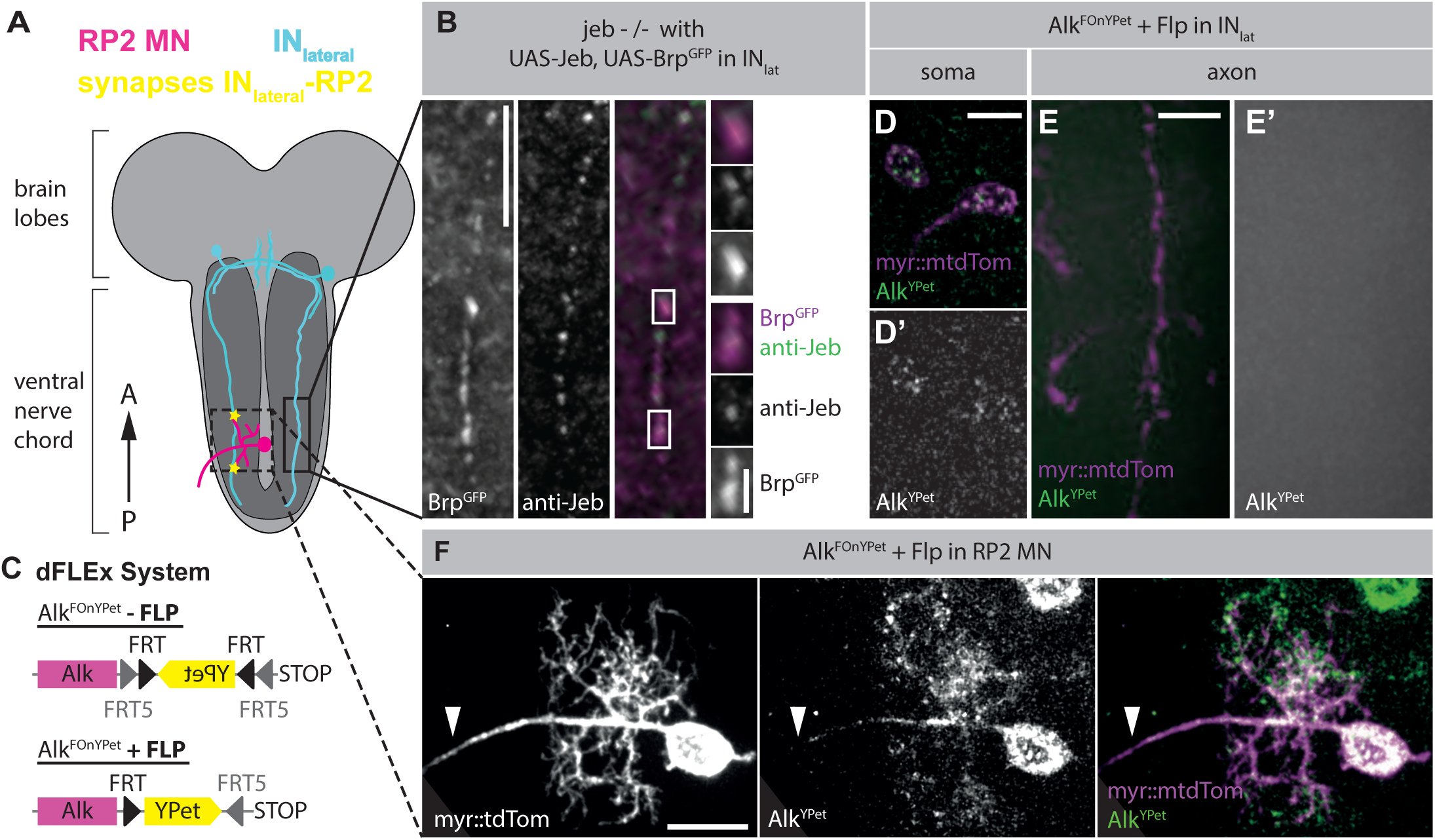
Subcellular localization of Jeb and Alk suggests synaptic anterograde signaling pathway. A – Schematic overview depicting the indentified pair of pre- (IN_lat_ interneuron, cyan) and post- synaptic (RP2 motorneuron, magenta) partner neuron in the ventral nerve cord (VNC) of *Drosophila* larvae; neuropil (dark grey), synaptic contact (astrerisk). B – Jeb (anti-Jeb) localizes to presynaptic release sites (BrpShort^GFP^) along IN_lat_ axons when Jeb expression is reinstated to IN_lat_ in *jeb* mutant background. C - Schematic of the conditional Alk^FOnYPet^ allele to label endogenous Alk expression. In cells that express FLP recombinase, a cassette between orthologous FRT and FRT5 will be stably inverted and thus leads to tagging of Alk at its endogenous genomic locus with a FLP dependent on-switchable YPet fluorophore (FOnYPet). Details in Fig S1 and STAR methods. D-E – Induced Alk^FOnYPet^ (D, green) reveals endogenous Alk^YPet^ localization to IN_lat_ soma (D, magenta); IN_lat_ axons are devoid of Alk^YPet^ (E, E’). F –Alk^YPet^ localizes to RP2 soma, primary neurite and dendrites, but does not enter the axon leaving the VNC (arrowhead). Scale bar = 2 µm (B, magnification), 10 µm (B, C, D), 20 µm (F)

We next aimed to reveal subcellular Alk localization. To this end we tagged Alk at its endogenous genomic 3’ end with an inducible YPet fluorophore, which is conditional on FLP expression (Alk^FOnYPet^, Fig 1C; details on dFLEx method see Fig S1H). We find that animals are homozygous viable and fertile for both, Alk^FOnYPet^ (tag not induced) and constitutively tag-induced Alk^YPet^, following induction in the parental germline. Because Alk loss of function is postembryonic lethal ^28^, we conclude that tagging endogenous Alk^FOnYPet^ leaves the protein functional. Induction of Alk^YPet^ in all neurons (via *nSyb-GAL4, UAS-Flp*) recapitulates an expression pattern previously identified by anti-Alk immunohistochemistry: Alk^YPet^ strongly localizes to the neuropil within the larval ventral nerve cord (VNC) and, in the adult brain, prominently labels the mushroom body calyx (Fig S1I-J) ^22, 23^. Using this method we asked if Alk is expressed in glia cells (using *repo-GAL4, UAS-FLP*) (Fig S1K). We were unable to detect Alk^YPet^, suggesting that, at least until the early third instar larval stage (48h after larval hatching (ALH)), Alk expression and, thus, Jeb-Alk signaling is exclusively neuronal.

When the Alk^YPet^ tag was induced in single RP2 motorneurons ^29^, we detected Alk^YPet^ fluorescence in the soma, primary neurite, and dendrites (Fig 1F), though not in axonal compartments (Fig 1F), nor at the neuromuscular junction (data not shown). Similarly, induction of the Alk^YPet^ tag in IN_lat_ (by *eyg-Gal4, UAS-Flp*) showed Alk^YPet^ localization in the soma, but not in axonal compartments (Fig 1 E,E’). These observations demonstrate that Alk protein localizes to postsynaptic, but not presynaptic, structures. Our protein localization data therefore imply that Jeb-Alk is an anterograde trans-synaptic signaling mechanism in central motor circuits, in agreement with previous findings in the visual system and the NMJ ^20, 21, 30^.

### Jeb regulates filopodial number but not presynaptic specializations

To determine the role of Jeb signaling in the CNS, we selectively removed Jeb function from selected cells using a genetically engineered conditional null allele. Briefly, we inserted an inducible artificial exon containing a translational stop codon and a transcription termination sequence into the endogenous Jeb locus (Fig 2A, methods), immediately preceding the region coding for the type A LDL receptor domain ^25^. The design of this construct is similar to the dFLEx approach outlined above, but uses Bxb1 integrase and corresponding attBx/attPx sites ^31^ (Fig 2A, details in Fig S2A). We term this technique BOnStop (Bxb1 dependent on-switchable stop codon). A similar, Flp recombinase dependent technique, called FlpStop, has recently been shown to be highly effective ^32^.

**Figure 2.**
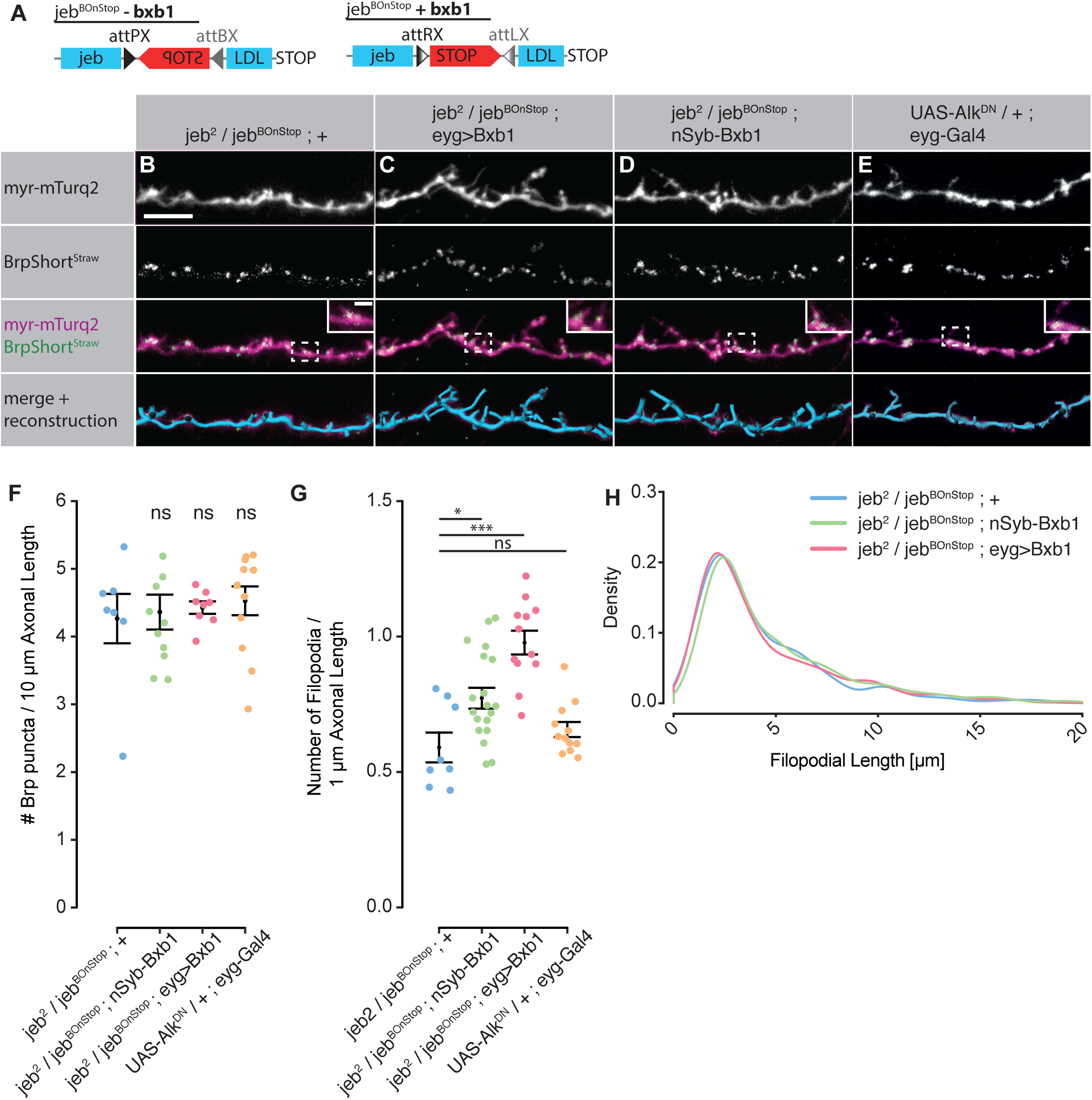
Presynaptic Jeb inhibits presynaptic filopodial growth via non-cell-autonomous feedback. A – Schematic of the conditional mutant jeb^BOnStop^ (Bxb1 dependent on-switchable STOP Codon) allele. Targeted Bxb1 integrase expression mutates Jeb in selected cells by induced inversion of an engineered exon to terminate translation and transcription immediately preceding the type A LDL receptor domain. For details see Figure 2S. B-E – (B) Control IN_lat_ axons (myr::mTurq2) at 48 h ALH are decorated with filopodial protrusions, many of which harbor a presynaptic specialization (BrpShort^straw^) at their insertion into the axon trunk (inserts). Presynaptic specialization were not found to enter filopodia. Targeted Jeb loss of function (C) exclusively in IN_lat_ and (D) in all neurons gives rise to increased filopodial growth, without obvious effects on numbers and localization of presynaptic specializations. (E) Inhibition of Alk signalling in IN_lat_ does not affect numbers of release sites, nor axonal morphology. F – Quantification of presynaptic specialization (BrpShort^Strawberry^) density along IN_lat_. G – Quantification of the numbers of filopodia along IN_lat_ axons. Normalized as numbers of filopodia per 1 µm axonal length. H– Density plot detailing the frequency of filopodial lengths reveals that overall filopodial morphology is unaffected by the loss of Jeb expression or inhibition of Alk. *p<0.05, **p<0.01, ***p<0.001, ns – not significant

We find that un-induced *jeb^BOnStop^ /jeb^2^* animals are fully viable and fertile, as would be expected for animals with one functional *jeb* allele. Embryos homozygous mutant for *jeb* fail to differentiate their visceral mesoderm, and therefore die during early larval stages ^24^. This mesodermal phenotype, with failure of gut formation, is reproduced following conversion of *jeb^BOnStop^* to the *jeb^Stop^* mutant allele by expression of Bxb1 integrase in the early mesoderm (*jeb^BOnStop^ / jeb^2^; Mef2>Bxb1*) (Fig S2B-C). This demonstrates that *jeb^BOnStop^* is a conditional *jeb* mutant allele, effectively induced in a cell and tissue-specific fashion by targeted Bxb1 integrase expression.

A previous report suggested that lack of Jeb-Alk function in the nervous system caused drastic locomotor defects, ultimately leading to death in late larval stages ^19^. However, when we induce Jeb loss-of-function in all neurons (by *jeb^BOnStop^ /jeb^2^*; *nSyb>Bxb1*), larvae survive until adult stages and do not show apparent locomotor deficits. Different roles have previously been attributed to Jeb signaling in the nervous system. At the larval NMJ, abrogation of synaptic Jeb anterograde signaling was reported to increase synaptic coupling strength, though without discernible structural change to presynaptic morphology ^19^. In the adult visual system, however, cell-autonomous knock-down of *jeb* in photoreceptor neurons caused a marked increase in the number of filopodia and their overgrowth into adjacent synaptic non-target layers ^20^.

Using our *jeb^BOnStop^* conditional loss of function allele, we sought to test the role of *jeb* in the larval VNC, specifically the locomotor network. Focusing on axonal and synaptic morphology of the identified IN_lat_ interneurons (Fig 2B-E), we find that both pan-neuronal (via *jeb^BOnStop^ / jeb^2^; nSyb>Bxb1,* 4.35 ± 0.26 Brp puncta/10µm, n=11) and cell-autonomous *jeb* loss of function (using *jeb^BOnStop^/jeb^2^; eyg>Bxb1,* 4.42 ± 0.09 Brp puncta/10µm, n=8) in IN_lat_ interneurons did not affect axonal presynaptic release site number over control (*jeb^BOnStop^ / jeb^2^; +,* 4.26 ± 0.42 Brp puncta/10µm, n=7, visualized using *BrpShort^straw^* ^33^) (Fig 2F), consistent with previous findings at the NMJ. However, as in photoreceptor axons, these *jeb* loss of function manipulations caused an increase in the number of filopodial side branches along the axon of the IN_lat_ interneuron (Fig 2G, control: 0.59 ± 0.05 filopodia/µm, n=8; pan-neuronal: 0.78 ± 0.04 filopodia/µm, n=12; cell-autonomous: 0.98 ± 0.04 filopodia/µm, n=19). Overall morphological characteristics of individual filopodia were not, however, altered (Fig 2H). We find that filopodia arise from bouton-like swellings, close to presynaptic release sites, but do not harbor markers indicative of either nascent presynaptic release sites (dSyd1::GFP; Fig S2F) or mature active zones (BrpShort^straw^; Fig 2B-E). This is in agreement with studies from developing tectal neurons in *Xenopus* embryos, where correlative live imaging-electron microscopy showed that such presynaptic filopodia do not contain immature nor mature synaptic release sites ^34^.

As might be predicted from the postsynaptic localization of the receptor for Jeb, Alk, cell-autonomous inhibition of Alk receptor function in this IN_lat_ interneuron (by *eyg-Gal4, UAS-Alk^DN^*) had no quantifiable impact on presynaptic axonal filopodia, confirming that Alk is not presynaptically active and arguing against autocrine signaling by Jeb through Alk (Fig 2G, 0.65 ± 0.03 filopodia/µm, n=12).

Presynaptic filopodia are known to be short-lived, highly dynamic structures ^1, 34^. To quantify their dynamic behavior, we developed intra-vital imaging methods for monitoring neuronal growth in the VNC (Fig 3). To immobilize larvae for the imaging period we adapted use of the anesthetic desflurane ^35^, and a piezo-electric motor to gently compress larvae between two cover slips in a controlled and reproducible fashion, thus improving optical contact and increasing imaging depth. Expression levels of a membrane marker in IN_lat_ allowed intra-vital imaging at 24 h ALH; the same animal was sacrificed 24 h later, at 48 h ALH, its VNC dissected and imaged immediately (Fig 3A-B). Following individual filopodia thus showed that only 34.5% (± 0.05%, n=4) persist from 24-48 h ALH in control animals (Fig 3C), and 70.0% (± 0.01%, n=4) of all filopodia emerge as new in this 24 h period (Fig 3D). Targeting cell-autonomous loss of *jeb* function to IN_lat_ did not significantly decrease overall filopodial stability (28.9 ± 0.01%, n=4, Fig 3C), but slightly increases the proportion of newly formed relative to stable filopodia at 48h ALH (77.8 ± 0.02%, n=4, Fig 3D). Taken together, our results, in agreement with previous data, show that Jeb down-regulates the formation of axonal filopodia with minor impact on individual filopodial dynamics, though it does not obviously influence general axonal morphology or numbers of release sites (Fig 2B-E).

**Figure 3.**
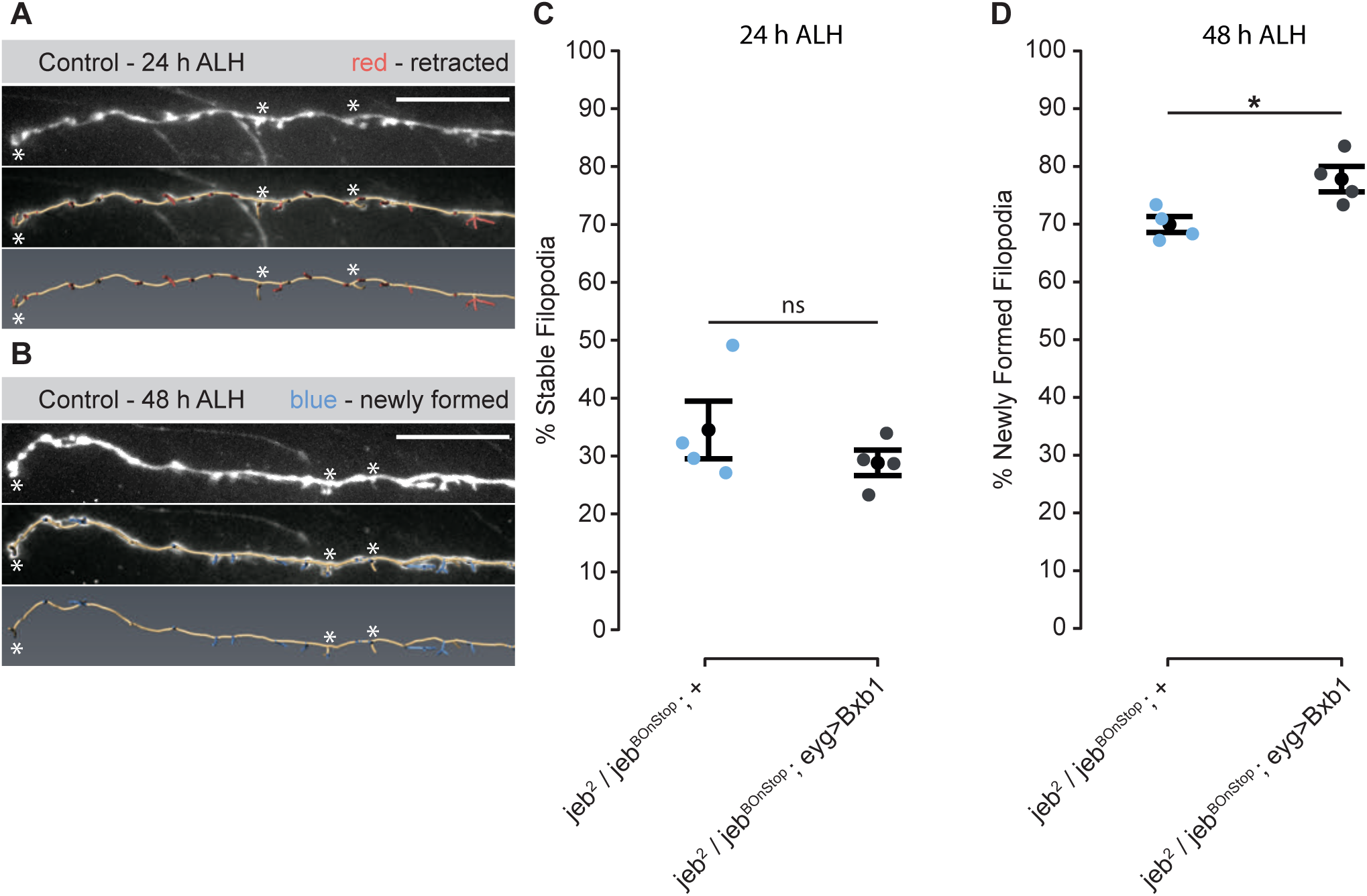
High structural turnover dynamics of INlat presynaptic filopodia are only slightly affected by loss of Jeb. A, B – Intra-vital recordings of INlat (myr::mtdTom) in (A) anesthetized larvae at 24 h and (B) in acutelty dissected nerve cords at 48 h ALH. Lower panels show an overlay with the 3D reconstruction of cell morphology. Red - Filopodia present at 24 h, and retracted until 48 h ALH; Blue - newly arising filopodia from 24 - 48 h. Asteriks - examples of persisting filopodia. C – Quantification of the proportion of filopodia removed from 24 to 48 h ALH (red in A). Loss of Jeb in INlat has no obvious effect on stability of individual filopodia in comparison to control. D - Quantification of the proportion of filopodia that newly form from 24 to 48 h ALH (blue in B). Loss of Jeb in INlat causes a slight increase in the proportion of newly formed filopodia in comparison to control. *p<0.05, ns – not significant

Exuberant growth of photoreceptor axons caused by loss of *jeb* has previously been interpreted as a defect in axonal targeting ^20^. However, we find that IN_lat_ axons successfully pathfind along the entire length of the *jeb* mutant VNC, suggesting that Jeb function is dispensable for guiding axonal growth, or spearheading the formation of presynaptic specializations. We therefore propose that presynaptic filopodia interact with postsynaptic dendrites of prospective partner neurons, as seen in *Xenopus* tectal neurons ^34^. Such inter-neuronal contact facilitates signaling by presynaptic Jeb activating dendritically localized Alk on the postsynaptic dendrite triggering a negative feedback signal that locally inhibits the formation of additional presynaptic filopodia, thus down-regulating locally further explorative filopodial behavior by the postsynaptic partner neuron.

### Alk signaling inhibits formation of postsynaptic specializations

Increased numbers of dynamic filopodia have been linked to synapse proliferation and synaptic plasticity in other systems ^3, 36^. Indeed, at the NMJ, loss of Jeb-Alk signaling causes strengthening of synaptic coupling ^19^. We therefore asked how changes in the number of presynaptic filopodia, that result from changes in Jeb-Alk signaling, might impact on synapse formation between central neurons (Fig 2F-H). To this end we established the first verified postsynaptic reporter for central neurons in *Drosophila*; DNA fragmentation factor related protein 2 (Drep2) has previously been suggested to be postsynaptically localized in Kenyon cells ^37, 38^. Using expansion microscopy (ExM, ^39, 40^ and a novel dFLEx^YPet^ allele for tagging the endogenous active zone protein Bruchpilot (Brp^FOnYPet^, Fig S3A), we could confirm that Drep2 profiles (anti-Drep2) in the VNC are juxtaposed to Brp^YPet^ marked presynaptic release sites. Interestingly, only a proportion of Brp^YPet^ puncta are opposed to anti-Drep2 marked postsynaptic sites, and we could determine that the Drep2-positive release sites are opposed to cholinergic release sites; acetylcholine being the principal excitatory neurotransmitter in insect nervous systems (visualized using *Brp^FOnYPet^; ChaT-T2A-GAL4, UAS-FLP*) (Fig S3B). In contrast, GABAergic release sites are not paired with Drep2 in the motor system (*Brp^FOnYPet^; GAD-T2A-GAL4, UAS-FLP*) (Fig S3C). Thus, our data show, that in the larval CNS, Drep2 principally localizes to cholinergic postsynaptic specializations.

Next, we quantified cholinergic postsynaptic specializations that form on individual RP2 motorneurons (RP2>FLP) by selectively inducing dFLEx^YPet^ tagging of endogenous Drep2 using a Drep2^FonYPet^ allele that we generated from a MiMIC site upstream of a *Drep2* coding intron, followed by induced homologous recombination (Fig S3D) ^41, 42^. Immunohistochemistry against the YPet fluorophore, followed by ExM, now allowed effective quantification of cholinergic postsynaptic specializations in RP2 motorneurons, as reported by Drep2^YPet^ profiles, opposed to Brp containing presynaptic release sites ^20^ (anti-GFP used to amplify YPet signal; Fig 4A-C). As expected for a postsynaptic protein, we find that Drep2 coincides with the RP2 membrane label (myr::tdTomato), but juxtaposes presynaptic Brp labelled sites at distance of 180nm (Fig 4D), in close agreement with previously reported distances for the Brp C-Terminus relative to the synaptic cleft ^43^. Importantly, we found that attempts of marking postsynaptic sites by overexpression of a GFP tagged version of Drep2 (*UAS-Drep2^GFP^*) in RP2 motorneurons causes abnormal dendrite development. This appears to be an over-expression artefact, which is circumvented when tagging the endogenous protein using Drep2^FOnYPet^ (Fig S3E-G, RN2FlpOut: 847.42 ± 27.19 µm; RP2>Drep2^FOnYPet^: 792.09 ± 18.56 µm, RP2>Drep2^GFP^: 669.95 ± 21.86 µm).

**Figure 4.**
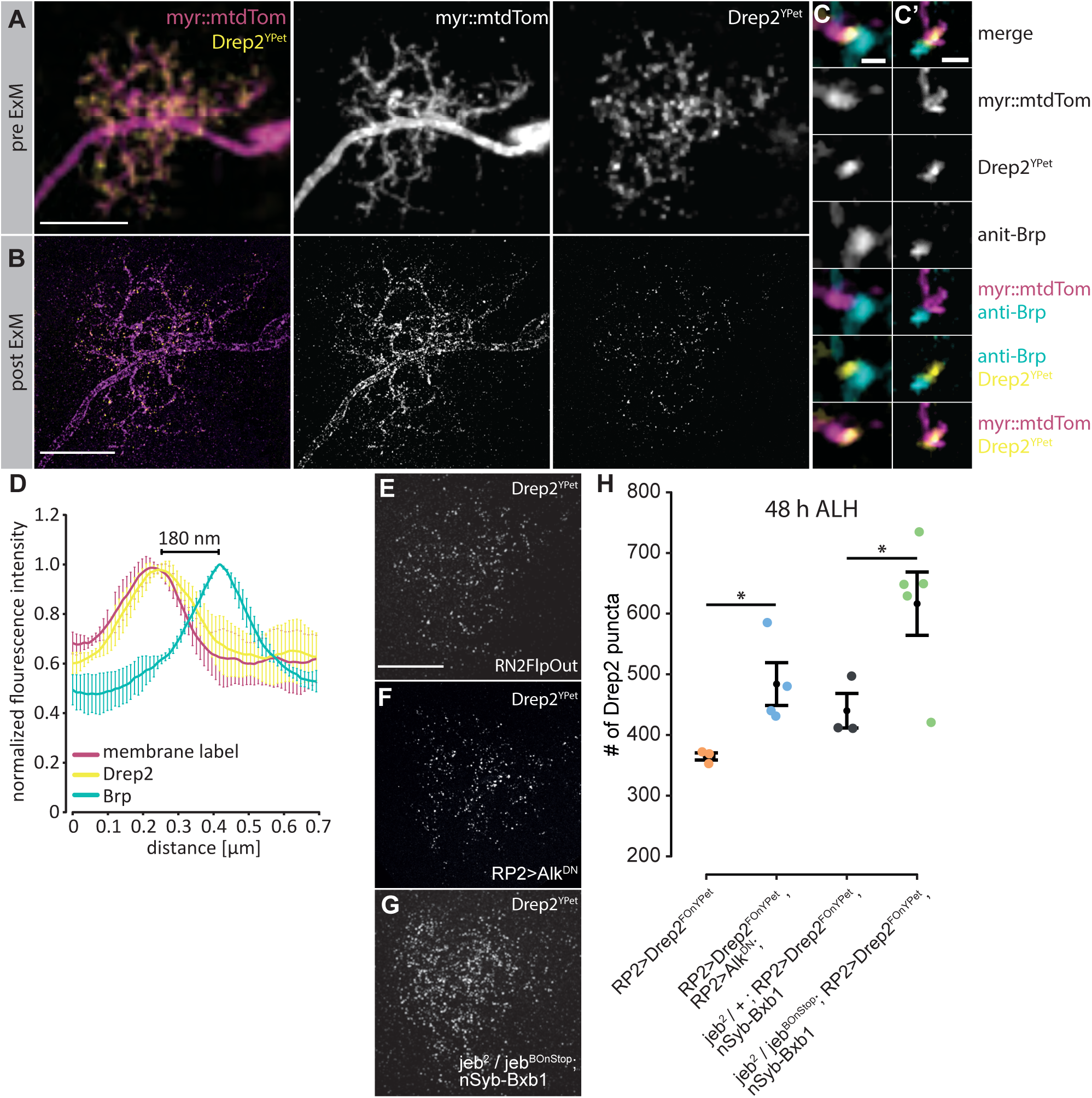
Jeb-Alk signaling inhibits formation of postsynaptic specializations. A –RP2 motorneuron (myr::mtdTom) with induced Drep2^FOnYPet^ imaged with confocal microscopy, demonstrating dendritic localization of Drep2^YPet^. B – The same RP2 motorneuron as in (A), now imaged with expansion microscopy (ExM). Individual postsynaptic specializations can now be distinguished and therefore accurately quantified. C-C’ - Partial maximum intensity z-projections showing representative synapses along dendritic branches in (B). Mature synapes are identified by apposition of postsynaptic Drep2^YPet^ (anti-GFP, yellow) with presynaptic Brp (anti-Brp, cyan). RP2 membrane is labeled with myr::mtdTom (magenta). D – Normalized mean intensity profile of Drep2^YPet^ (yellow), Brp (cyan) and dendritic membrane label (myr::mtdTom, magenta) across synaptic contacts. Drep2^YPet^ localization intimately overlaps with the dendritic membrane. The distance between mean intensity maxima of Drep2^YPet^ and the cytosolic terminal (C-Term) of Brp was measured as 180nm. n=15 synapses from 3 RP2 dendrites. E-G – Max. intensity z-projections of endogenous Drep2^YPet^ in RP2 dendrites after ExM in (E) control, (F) pan-neuronal jeb mutant, and (G) single cell Alk inhibition. H – Quantification of the number Drep2^Ypet^ apposed with Brp along RP2 dendrites at 48 h ALH demonstrates that Jeb-Alk signaling inhibits formation of postsynaptic specializations. Welch two-sample t-test *p<0.05. Scale bar = 10 µm (A,B,E) 500 nm (C)

We find that cell-autonomous inhibition of Alk in RP2 motorneurons, through expression of a well characterized dominant negative acting form (*UAS-Alk^DN^* ^20^), causes a notable 1.3-fold increase (RP2>Alk^DN^: 484.00 ± 30.58, n=4) of Drep2 puncta as compared to control (364.67 ± 4.82, n=3, Fig 4F,H). When Jeb-Alk signaling is removed throughout the nervous system (*jeb^BOnStop^/jeb^2^; nSyb-Bxb1: 616.40* ± 46.68, n=5; Fig 4G,H) this increase in postsynaptic specializations is further enhanced 1.4 fold (*+/jeb^2^; nSyb-Bxb1:* : 440.00 ± 23.27, n=3; Fig. 4G,H). Interestingly, these increases in postsynaptic sites in RP2 motorneurons, following cell-autonomous Alk inhibition, only manifest during postembryonic life, and are not evident at hatching (0 h ALH, Fig S3H, control: 72.67 ± 4.81; RP2>Alk^DN^: 73.67 ± 4.48).

It is reasonable to suppose that an increase in postsynaptic specializations might require a concomitant increase in presynaptic sites. However in *Drosophila*, most central synapses are polyadic, with multiple postsynaptic partners opposed to a single presynaptic specialization ^44–46^. Because we do not find loss of Jeb-Alk signaling impacting the number of presynaptic specializations (Fig 2F), yet leading to increases in postsynaptic sites within motorneuron dendrites (Fig 4H), our data suggest that this signaling pathway might modulate the number of postsynaptic specializations that pair up with any one presynaptic release site.

In mouse pyramidal neurons synaptogenesis appears competitive, such that less tenacious synapses will be withdrawn in favor of stronger ones, and neurons that outcompete others proliferate more synapses and grow a bigger dendrite ^47^. Here, we showed that Alk signaling deficient motorneurons markedly increase the numbers of synaptic input sites they form, irrespective of whether just one, or all neurons (i.e. pan-neuronal) lost Alk signaling. This suggests that Alk does not influence competitive performance, but that proliferation of postsynaptic sites is regulated cell-autonomously by Alk activation.

### Jeb-Alk signaling inhibits developmental strengthening of synaptic excitation and stabilizes circuit function

Increased synaptic connectivity, as seen at the anatomical level, suggests a corresponding increase in synaptic coupling strength. To test this directly, we performed intracellular recordings from RP2 motorneurons to quantify rhythmic cholinergic excitatory synaptic inputs that occur during fictive crawling activity. These excitatory currents are termed spontaneous rhythmic currents (SRC) ^48^. Indeed, we find that SRCs recorded in RP2, in Jeb-/- show significantly increased duration indicative of heightened synaptic coupling (Fig 5A-C; *jeb^BOnStop^/jeb^2^; +*: 0.51 ± 0.04 s, n=10; *jeb^BOnStop^/jeb^2^; nSyb-Bxb1:* 1.15 ± 0.06 s, n = 10). SRC frequency (Fig 5D, *jeb^BOnStop^/jeb^2^; +*: 15.83 ± 2.77 min^-1^, n=10; *jeb^BOnStop^/jeb^2^; nSyb-Bxb1:* 12.23 ± 3.02 min^-1^, n=10) and SRC current density (Fig 5E, *jeb^BOnStop^/jeb^2^; +*: 35.14 ± 3.30 pA/pF, n=10; *jeb^BOnStop^/jeb^2^; nSyb-Bxb1:* 43.07 ± 2.69 pA/pF, n=10) are not significantly affected. This outcome is consistent with increased numbers of postsynaptic specializations, that we also observe in a Jeb-/- nervous systems. Increased SRC duration is a characteristic of *Drosophila* models for epilepsy ^48^. Such larvae exhibit prolonged duration of seizure recovery following electroshock. We find that seizure recovery time approximately doubles in pan-neuronal Jeb^-/-^ larvae at 48 h ALH compared to control (Fig 5F, *jeb^BOnStop^/jeb^2^; +*: 107.2 ± 7.72 s, n=30; *jeb^BOnStop^/jeb^2^; nSyb-Bxb1:* 212.9 ± 11.13 s, n=30). By contrast, removal of Jeb function from all neurons does not significantly affect synaptic excitation (SRC duration) at larval hatching (0h ALH, Fig 5G-K), in line with Jeb-Alk signaling not affecting numbers of postsynaptic specializations during embryonic development (Fig 4G-H, Fig S3H).

**Figure 5.**
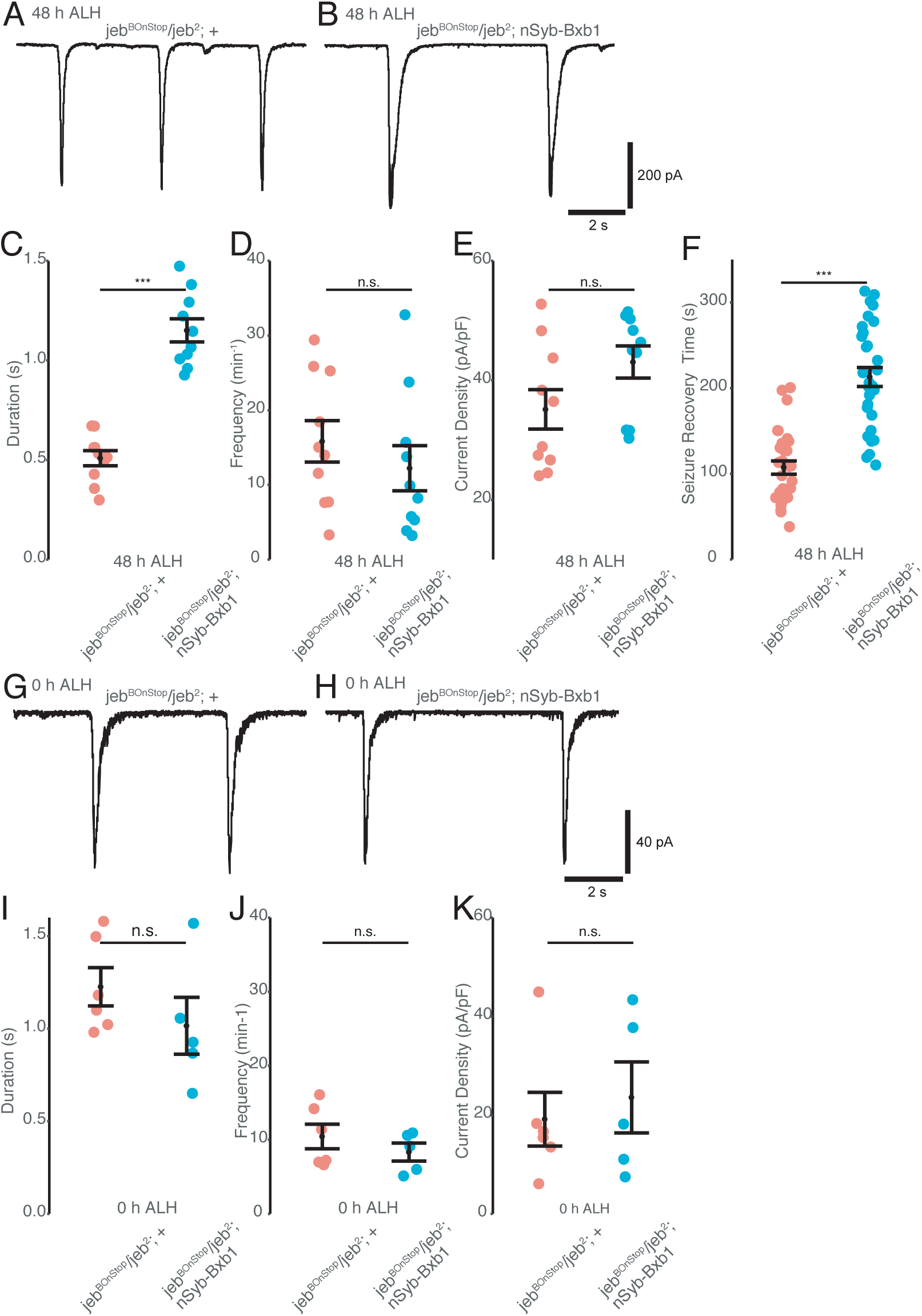
Pan-neuronal loss of Jeb results in an developmental increase of synaptic excitation and epilespy-like seizure phenotype. A-E – Whole-cell patch recordings of SRCs from identified RP2 motoneurons in dissected larvae at 48h ALH, from control (A) and pan-neuronally Jeb-/- (B) animals. Loss of Jeb causes strongly increased synaptic excitation at 48h ALH evident by increased SRC duration (C), without significant changes to SRC frequency (D) and current density (E). F – Seizure recovery times after electro-shock are doubled in pan-neuronally Jeb-/- larvae at 48h ALH larvae in comparison to control. G-K –SRCs recorded at 0h ALH from RP2 motorneurons are indistinguishable between control (G) and pan-neuronal Jeb-/- (H) larvae: SRC duration (I), frequency (J) and current density (K) are not significantly different. ***p<0.001, ns – not significant

We conclude that the developmental over-proliferation of cholinergic postsynaptic specializations, as seen at the anatomical level, in Jeb-Alk deficient nervous systems (Fig 4) gives rise to increased synaptic excitation to RP2 motor neurons, and results in a motor circuit with decreased ability to recover from seizure. Thus, Jeb-Alk signaling is required to promote stability of circuit function and resilience.

### Cell-autonomous and pan-neuronal Alk inhibition result in distinct dendritic growth phenotypes

Dendrites are the substrate on which postsynaptic specializations form. Therefore, synapse proliferation and dendritic structural elaboration are often thought of as intimately linked ^49^. Indeed, synaptogenesis can stabilize nascent axonal and dendritic branches and thus promote structural expansion of synaptic terminals ^1, 2^ (synaptotropic growth, reviewed in ^50^). In the case of RP2 motorneurons, as the animal grows, their dendrites enlarge and proportionately increase the number of supported presynaptic inputs ^14, 15^. However, there are observations that question a strict co-regulation of dendritic and synaptic growth. For example, during network formation, experimentally increased availability of presynaptic sites actually results in reduced dendritic arbors, suggesting that synapse formation can also inhibit dendritic growth, at least during early developmental stages ^51^.

To test the interdependency between synapse formation and dendritic elaboration, we first analyzed dendritic arbor morphology of RP2 motorneurons (Fig 1A yellow) in pan-neuronal *jeb^-/-^* mutant animals (*jeb^BOnStop^ /jeb^2^; nSyb-Bxb1*). We found that total dendritic length (TDL) of RP2 motorneurons was significantly increased in pan-neuronal *jeb^-/-^* mutant animals (1099.49 ± 26.20 µm) compared to control (959.91 ± 12.56 µm, Fig 6A-F), in line with the synaptotropic growth hypothesis ^52^. Gross morphology and targeting of dendrites to the characteristic RP2 territory were normal, indicating that dendritic pathfinding was unaffected. Pan-neuronal inhibition of Alk (*elav-Gal4, UAS-Alk^DN^*) caused a comparable increase in TDL (Fig 6A-F, 1193.23 ± 79.50 µm), confirming that Alk is the only Jeb receptor, at least in neurons. Thus, pan-neuronal loss of Jeb-Alk signaling induces a strong increase in presynaptic filopodia (Fig 2G), increased overall length of postsynaptic dendritic arbors and increased postsynaptic connectivity. The number of presynaptic specializations, however, remains unaffected.

**Figure 6.**
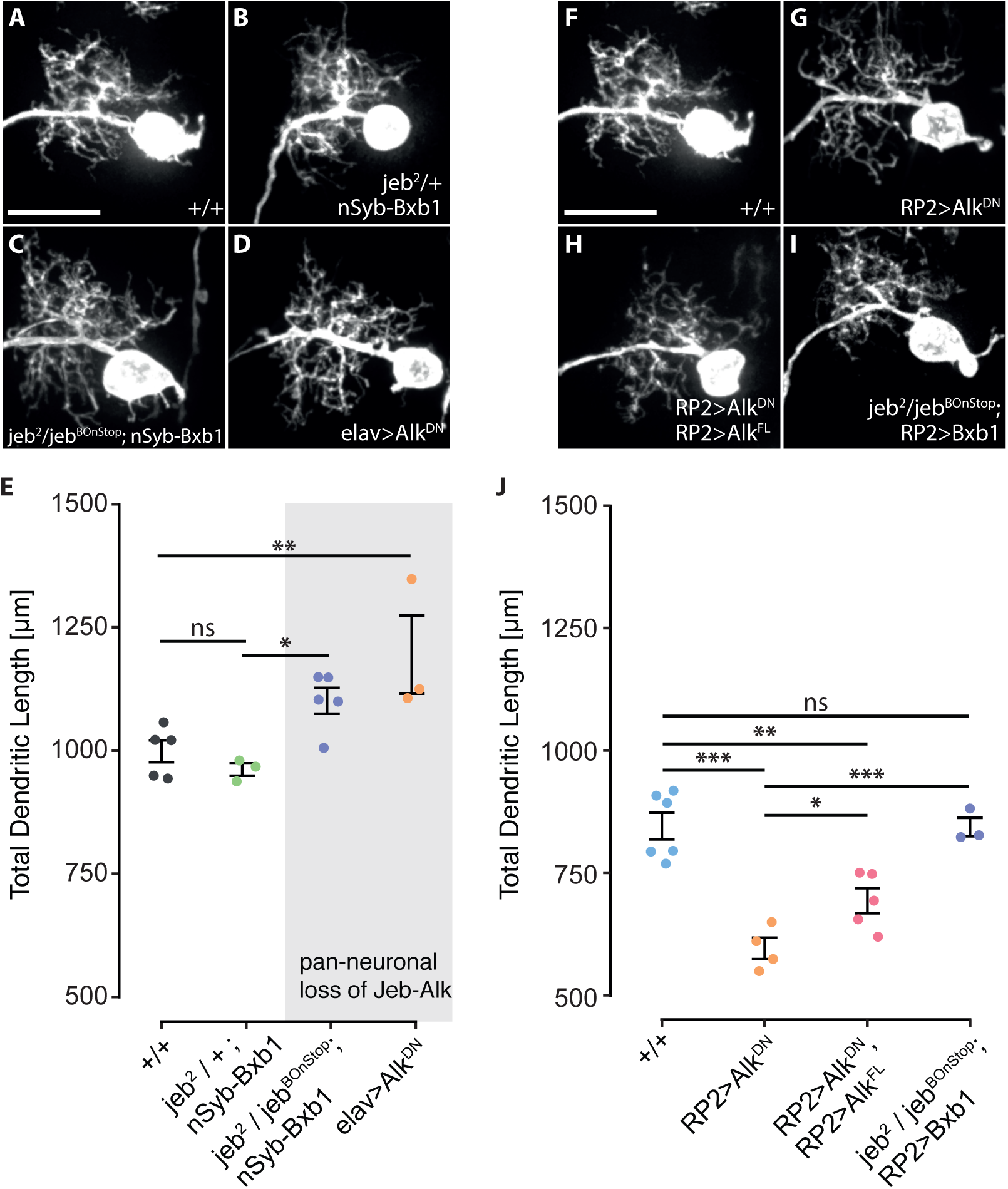
Cell-autonomous and pan-neuronal abrogation of Jeb-Alk signalling results in distinct growth phenotypes of RP2 motorneuron dendrites. A-E – Dendrites (RP2>myr::YPet) grow larger under pan-neuronal abrogation of Jeb-Alk signalling. (A,B) control, (C) pan-neuronal jeb mutant (nSyb>Jeb^BOnStop^), and (D) pan-neuronal Alk inhibition (Elav> Alk^DN^). (E) Quantification of RP2 total dendritic length in genotypes shown in A-D. F-J – Dendritic growth is reduced upon cell-autonomous inhibition of Alk signalling in RP2 only. (F) control, (G) cell-autonomous Alk inhibition in RP2 motorneuron (RP2>Alk^DN^), (H) co-overexpression of Alk^DN^ and full-length Alk (Alk^FL^) in RP2, and (I) induced jeb mutation in RP2 only. (J) Quantification of RP2 total dendritic length in genotypes shown in (F-I). *p<0.05, **p<0.01, ***p<0.001, ns – not significant

The increased dendritic length in Jeb-Alk deficient nervous systems might be a cell-autonomous consequence of the lack of postsynaptic Alk activation. Alternatively, this postsynaptic overgrowth might be regulated at the tissue level. To discriminate between these two possibilities, we targeted Alk^DN^ expression to single RP2 motorneurons (Fig 6G-I). We found that inhibition of Alk in single cells caused a significant reduction of RP2 TDL as compared to control (control: 847.42 ± 27.19 µm, n=6; RP2>AlkDN: 589.09 ± 24.84 µm, n=4). This stunted dendritic growth phenotype could be partially rescued by co-expression of full length Alk (Fig 3I+K, 695.25 ± 25.56 µm), confirming the specificity of Alk^DN^ for Alk signaling. This observation shows that Jeb-Alk signaling cell-autonomously promotes dendritic growth; however, global abrogation of Alk activation at the tissue-level changes intercellular interactions toward stimulating RP2 motorneuron growth. Throughout, Jeb acts exclusively in anterograde signaling, further underlined by the fact that cell-autonomous knockdown of *jeb* in RP2 motorneurons (*jeb^BOnStop^ / jeb^2^; RP2>Bxb1, 845.51* ± 18.85 µm) does not impact on normal dendritic growth (Fig 6J).

### Jeb-Alk signaling regulates elaboration of dendritic arbors exclusively during postembryonic circuit growth

We showed above that experimental Jeb-Alk inhibition non-cell autonomously leads to an increase in presynaptic filopodia, suggesting that normally Alk activation by Jeb acts to locally reduce the number of presynaptic filopodia (Fig 2G). Filopodial protrusions can form pioneering contacts with neighboring cells, thus speeding up target discovery and promoting synaptogenesis between partners ^9, 34, 36^. Consequently, we consider that interactions between presynaptic filopodia and nascent postsynaptic dendritic branches are capable of promoting dendritic arbor elaboration. Widespread formation of super-numerous presynaptic filopodia, as seen with pan-neuronal loss of *jeb*, would thus not only result in increased dendritic complexity, as we have observed, but also heightened growth dynamics. In contrast, Alk inhibition in individual RP2 motorneurons is unlikely to cause effects on overall presynaptic filopodial dynamics and therefore is similarly unlikely to result in changes to dendritic growth dynamics. However, cell autonomous inhibition of Alk activity in single motorneurons leads to significantly more excitatory cholinergic inputs forming on these, which might in turn result in reduced dendritic growth, as previously shown ^51^.

To test the validity of this model we set out to quantify the dynamics of dendritic growth by intra-vital microscopy of individual RP2 dendritic arbors in transiently anaesthetized larvae immediately after hatching, at 0 h ALH, followed up in 24 h intervals, at 24 and 48 h ALH (early second and third instar stages, respectively; the last time point in acutely dissected nerve cords, Fig 7A). We find that RP2 dendrites double in size every 24 h, consistent with previous findings ^15^.

Relative to controls, not subjected to intra-vital imaging and recorded at 48 h ALH, we observed a slight, but non-significant reduction in total dendritic length following this intra-vital imaging sequence (<5%, Fig 7B, control: 847.42 ± 27.19 µm; anesthetized: 807.95 ± 36.52 µm). We therefore conclude that larvae subjected to repeated anesthesia display normal dendritic development. Importantly, at the time of larval hatching (0 h ALH), RP2 dendritic arbor size did not differ between control condition (184.99 ± 6.77 µm), cell autonomous inhibition of Alk (193.48 ± 8.29 µm) or pan-neuronal loss of *jeb* function (186.25 ± 4.58 µm) (Fig 7C). However, during subsequent postembryonic stages, cell autonomous inhibition of Alk leads to smaller RP2 dendritic arbors, 17% reduction in TDL at 24 h ALH (control: 404.40 ± 23.58 µm, RP2>Alk^DN^: 334.91 ± 16.35 µm), and this difference is maintained at 17% until 48 h ALH (control: 807.95 ± 36.52 µm, RP2>Alk^DN^: 673.81 ± 42.774 µm). Similarly, pan-neuronal loss of *jeb* causes a slight though statistically not significant increase in RP2 dendritic length at 24 h ALH (8%, 435.2 ± 20.77 µm); this then propagates to a significant 31% increased TDL by 48 h ALH (1056.91 ± 69.18 µm).

This shows that neuronal Jeb-Alk signaling does not regulate dendritic growth during initial circuit formation in the embryo, when activity dependent mechanisms likely play a dominant role in shaping nascent motor circuitry ^48, 53^; but it is during post-embryonic larval stages that Jeb-Alk signaling regulates the rate of dendritic growth.

**Figure 7.**
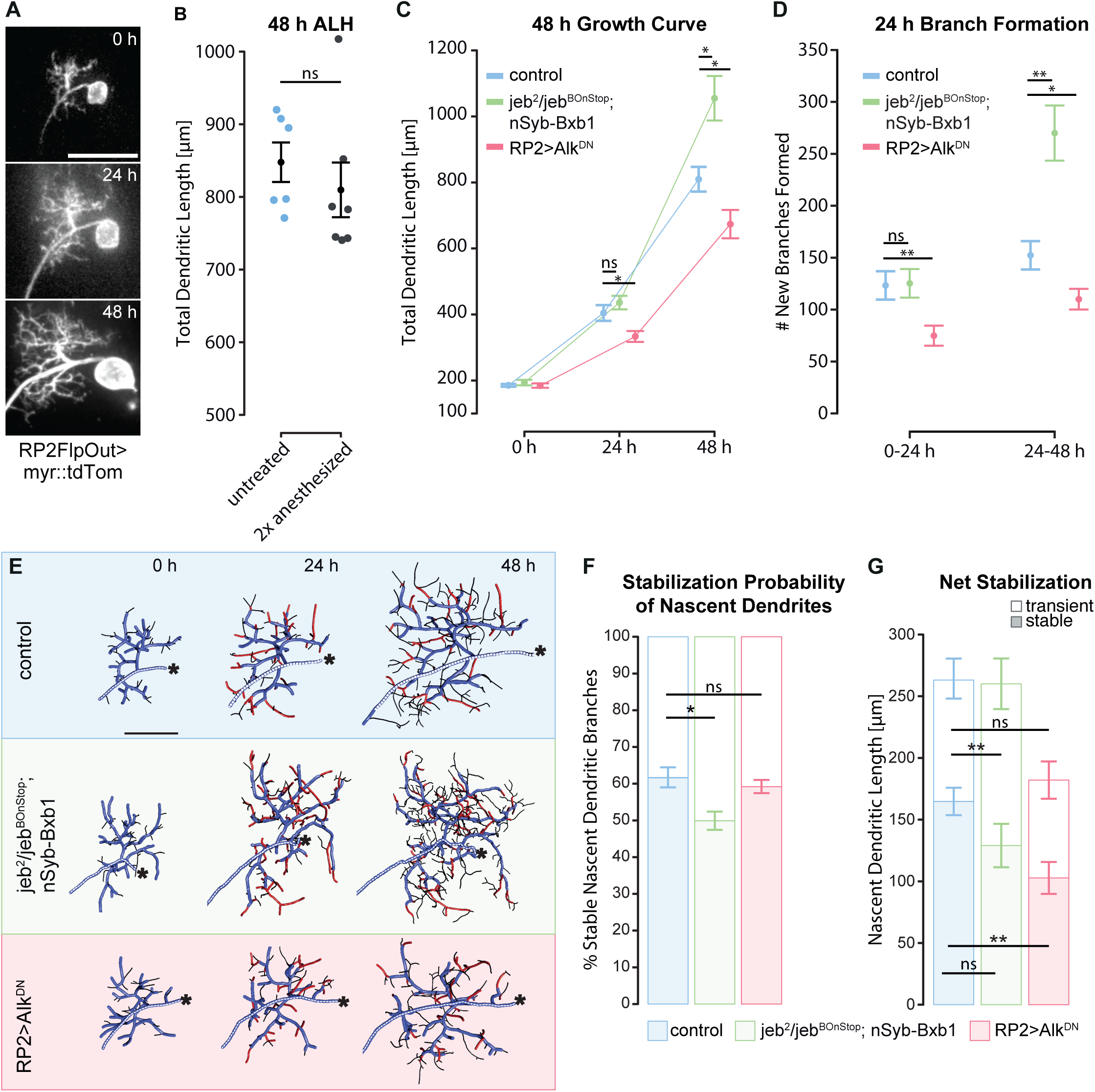
RP2 dendritic turnover dynamics scale with presynaptic filopodia dynamics. A – Intra-vital imaging of RP2 dendrites at 24 h intervals. Recordings at 0 an 24 h ALH were acquired from anesthetized larvae. At 48 h ALH, the VNC was dissected and acutely imaged. Scale bar – 20 µm B – Quantification of total dendritic length shows that dendrite development is not significantly affected by twice repeated anesthetization during intra-vital imaging in comparison to untreated control. C – Quantification of total dendritic length from intra-vital recordings reveals that Jeb-Alk signalling regulates progressive dendritic growth exclusively during postembryonic stages. RP2 cell-autonomous inhibition of Alk (red) slows postembryonic dendritic growth; pan-neuronal loss of Jeb (green) causes progressively increasing dendritic growth in comparison to control (blue). Embryonic dendritic development is not regulated by Jeb-Alk, evident by the lack of obvious dendritic phenotypes at 0 h ALH. D – Quantification of the formation of new dendritic branches shows that RP2 cell-autonomous inhibition of Alk signalling (red) slows progressive dendritic growth due to a reduced rate of branch formation. In contrary, pan-neuronal abrogation of Jeb (nSyb>Jeb^BOnStop^) promotes branch formation from 24 – 48 h ALH in comparison to control. E – Reconstructions of RP2 dendrites, colour-coded for branch lengths that were a) stable over all three timepoints (blue); b) newly formed from 0 to 24 h ALH, and stabilized until 48 h ALH (red); c) filopodia not covered by a) and b) (black). Asterisk - position of cell body. F – Quantification of the stabilization probability of nascent dendritic branches. The percentage of branches that newly formed from 0 to 24 h ALH, and subsequently stabilized until 48 h ALH, is significantly reduced under pan-neuronal loss of Jeb, but not affected by RP2 cell-autonomous inhibition of Alk in comparison to control. G – Quantification of the net addition of dendritic length. Stacked bar plots show the summed length of dendritic branches that formed from 0 to 24 h ALH and a) persisted until 48 h ALH (stable length), or b) subsequently retracted (transient length). Dendrites in pan-neuronal jeb mutant background display reduced net growth despite an increased branch formation from 24 – 48 h ALH. Blue – control (RP2>myr::mtdTom); Green - pan-neuronal loss of Jeb (nSyb>Jeb^BOnStop^); red – RP2 cell-autonomous inhibition of Alk (RP2>Alk^DN^). *p<0.05, **p<0.01, ns – not significant

### Changes in presynaptic filopodial dynamics are paralleled by changes in dendritic growth behavior

To enable net growth, stabilization of nascent neuronal branches must be greater than branch retraction rates. To quantify the dynamics that give rise to dendritic arbor growth, we tracked individual dendritic branches from 0 h to 48 h ALH and determined rates of branch formation, retraction and stabilization (Fig 7D-G). New branches are added throughout the dendritic tree (Fig 7D-E, 0-24 h: 134.14 ± 14.43, n=7; 24-48 h: 177.00 ± 15.25, n=7). Importantly, 61.74 ± 2.72 % of branches formed from 0h to 24h ALH persist until 48h ALH in control larvae (Fig 7F). This shows that nascent dendritic branches are considerably more stable than presynaptic filopodia of IN_lat_, of which only 34.5% are maintained during this period (Fig 3).

We next tested the dendritic growth dynamics in animals with pan-neuronal loss of *jeb*. Here, RP2 dendrites show a markedly elevated rate of branch formation (Fig 7D; 24-48 h: 305.75 ± 27.61 branches, n=4) and at the same time reduced stabilization probability (49.95 ± 2.49%, n=4), as compared to dendrites from controls (Fig 7F). Despite this increased rate of branch formation, the total length of stable dendritic branches, that had formed in the first 24 hours of larval live (0-24 h ALH) and persisted for the next 24 hours (until 48 h ALH), is slightly shorter than in control (Fig 7G, control: 164.47 ± 11.13 µm, n=7; pan-neuronal *jeb* mutant: 128.78 ± 11.13 µm, n=4). Thus, stability of RP2 dendrites is reduced in a nervous system mutant for *jeb*. In contrast, cell autonomous inhibition of Alk, in single RP2 motorneurons, reduced formation of new branches (Fig 7D, 0-24h: 86.43 ± 10.14 branches, n=7; 24-48 h: 131.71 ± 11.80 branches, n=7), without significant effect on their stabilization probability (Fig 7F, 59.24 ± 1.80%, n=7).

In conclusion, we find that dendritic branch formation and retraction dynamics scale with overall presynaptic filopodial dynamics, and that these are not cell-autonomously regulated by Alk signaling. Formation of postsynaptic sites, in turn, likely acts inhibitory on the formation of new dendritic branches, possibly by reducing sensitivity to interactions with presynaptic filopodia.

## Discussion

Animals need to maintain neuronal circuit function while adapting to a body growing in size or changing environmental conditions. Mechanisms that regulate the initial assembly of neuronal circuits have been studied intensively. However, we know much less about how growth is regulated during post-embryonic phases, following the integration of neurons into functional circuits. Here we show that trans-synaptic Jeb-Alk signaling acts to slow motor circuit growth upon synapse formation in *Drosophila* larvae, and as a result, maintains functional stability during postembryonic growth. Jeb is released from cholinergic terminals and activates postsynaptic Alk, which negatively regulates the proliferation of post-, but not pre-, synaptic specializations. Alk activation by Jeb also seems to elicit an as yet uncharacterized retrograde signal that inhibits the formation of presynaptic filopodia. We find that dendritic structural plasticity is promoted by presynaptic filopodia, but repressed by increasing numbers of postsynaptic specializations.

Based on our observations, we propose a model for how postembryonic growth is coordinated between synaptic partner neurons, namely that presynaptic filopodia enhance dendritic growth. Once a dendritic protrusion contacts a presynaptic release site, presynaptic Jeb induces dendritic Alk activation. This Alk activation then acts to inhibit addition or differentiation of further postsynaptic specializations nearby. It also elicits retrograde signaling that dampens presynaptic filopodial dynamics, thus curbing presynaptic exploratory contacts in the immediate vicinity. Our data point to Alk-receptor activation being interpreted locally at a forming synaptic contact, therefore repressing nearby formation of other contacts and thus reducing the density of postsynaptic specializations along a dendritic arbor.

### Distinct mechanisms control initial circuit assembly and later growth

Strengthening of synaptic drive to motorneurons, during development, is achieved by increasing the number of synaptic connections (this study, also ^14, 15^). The mechanisms that maintain network function during postembryonic life are likely different from those that allow function to emerge initially during an identified early critical period of heightened activity-regulated plasticity ^48, 54, 55^. These emergent network configurations cannot be overruled or reprogrammed through later interference with network activity ^54^. Here, we show that Jeb-Alk signaling impacts on dendritic elaboration and connectivity, but only during the post-embryonic growth phase, where it acts to maintain stability of circuit function. The nature of the switch from initial network assembly mechanisms to subsequent programs of maintenance and stability during periods of growth and expansion is not known. One candidate is developmentally and nutritionally regulated steroid hormone signaling, which we previously showed gates postembryonic neuronal growth ^15^. Future work targeting the upstream mechanisms of Alk and steroid signaling might reveal possible therapeutic targets to lift the rigidity of circuit maintenance, and re-instate embryonic flexibility to support recovery from, for example, brain damage or neurodegenerative disorders.

### Equilibrated circuit growth to maintain circuit function

During postembryonic circuit growth, synaptic partners expand their neuronal arbors and synaptic sites by equilibrated addition of pre- and postsynaptic specializations, thus preserving embryonically established patterns of synaptic connectivity throughout postembryonic circuit growth ^14, 16^. Therefore, equilibrated expansion of such synaptic patterns likely underlies the maintenance of circuit function during organismal growth ^56^. Reciprocal growth promotion alone is sufficient to equilibrate cell proliferation of a two-cell network. Negative feedback stabilizes this reciprocal promotion and thus prevents uncontrolled over-proliferation ^57^. Contrary to this ^19^, we find that Jeb mutant nervous systems support normal crawling performance. This demonstrates that overall network function is maintained despite massive over-proliferation of cholinergic postsynaptic specializations in a nervous system entirely mutant for Jeb-Alk signaling (Fig 3). In the same vein, loss of Alk function in adult *Drosophila* mushroom body increases memory performance, demonstrating that learning circuits are not impaired, by contrast performance is increased ^23^. We thus speculate that Jeb-Alk signaling does not severely impact synaptic partner specificity between neurons within a circuit. However, continuous over-growth of neuronal circuits is space- and energy consuming, and we show that it leads to increased cholinergic excitation paralleled with reduced functional resilience, as manifested by significantly prolonged durations required for recovery from electro-shock induced seizures, Fig 5. We therefore suggest that Jeb-Alk signaling might operate to keep positive growth interactions at bay, and thus act to maintain and stabilize embryonically established circuit function possibly as a negative feedback signal to synapse formation.

### Presynaptic filopodia evoke postsynaptic structural plasticity

At the developmental stage when neuronal projections have arrived in common meeting regions and synaptogenesis commences, axonal and dendritic filopodia change their role from steering neurite outgrowth, to negotiating synapse placement ^4, 12, 58, 59^. Increased filopodial dynamics correlate with phases of heightened synaptogenesis during postembryonic development ^3^, and factors that strengthen synaptic coupling also induce an increase in filopodia formation ^60, 61^. These data suggest that filopodial activity may be crucial to drive circuit plasticity and synaptogenesis. Studies in dissociated neuron culture revealed that axonal filopodia frequently contain presynaptic proteins and synaptic vesicles, and are able to quickly assemble presynaptic specializations upon contacting a target cell ^5, 6^. Similarly, dendritic filopodia can induce presynaptic differentiation ^4^, then stabilize to transform into postsynaptic spines ^12^. In intact *Xenopus* tectum, however, the majority of presynaptic release sites emerge on stable axonal branches, whereas postsynaptic specializations readily form within dynamic dendritic branches ^34^. Our data from *Drosophila* postembryonic motor circuit growth support the latter scenario: 1) dynamic IN_lat_ axonal filopodia are empty of both early and late molecular markers of presynaptic differentiation (dSyd1, and Brp respectively); and 2) in a *jeb-/-* mutant nervous system, supernumerary postsynaptic specializations form within motorneuron dendrites despite significantly increased dendritic turnover. We therefore suggest that axonal and dendritic filopodia play conceptually different roles during circuit growth: presynaptic filopodia induce sprouting of dendritic filopodia and thus attract dendritic growth towards presynaptic specializations; subsequently, postsynaptic specializations form.

Taken together, our findings suggest fundamentally different behaviors for pre- *versus* postsynaptic specializations during postembryonic circuit growth: proliferation of presynaptic specializations is fairly static, while postsynaptic specializations readily form and retract. This could explain why, in this system, the number of presynaptic release sites of the IN_lat_ axons shows much less variability (Fig 2) than connectivity patterns with identified postsynaptic partners ^14^.

### Dendritic shape, synapse placement and behavioral output

Dendrites, due to their geometry, are inherently endowed with mechanisms for weighting and integrating synaptic input depending on its localization along the dendritic arbor. Thus dendritic shape and synapse placement can have a strong influence on neuronal computational performance (reviewed in ^62, 63^). However, recent studies in the stomatogastric ganglion of crabs showed that the specific tapering of dendritic branches can effectively compact elaborate dendritic structures to a uniform electrotonic compartment, thus strongly reducing the impact of synapse positioning along a dendritic arbor on signal integration and ultimately spike output ^64–66^. Conversely, experimentally reduced dendritic arbor complexity of *Drosophila* flight motorneurons still supports astonishingly normal neuronal performance, and only behaviors requiring fine precision appear to be negatively affected ^67^. Here we show that a drastic increase in dendritic arbor size, and numbers of synaptic connections, as caused by pan-neuronal abrogation of Jeb-Alk signaling (Fig 4 and 5) did not produce noticeable motor defects. This demonstrates that also in the *Drosophila* larval motor system, the fine details of dendritic morphology may not be of significance for the computation of adequate neuronal output. In conclusion, we propose that proportional growth of dendrites leads to a proportional strengthening of existing synaptic patterns; whereas changes in the positioning of dendritic subtrees would alter the choice of synaptic partner neurons and thus also the balance between those.

## Online Methods

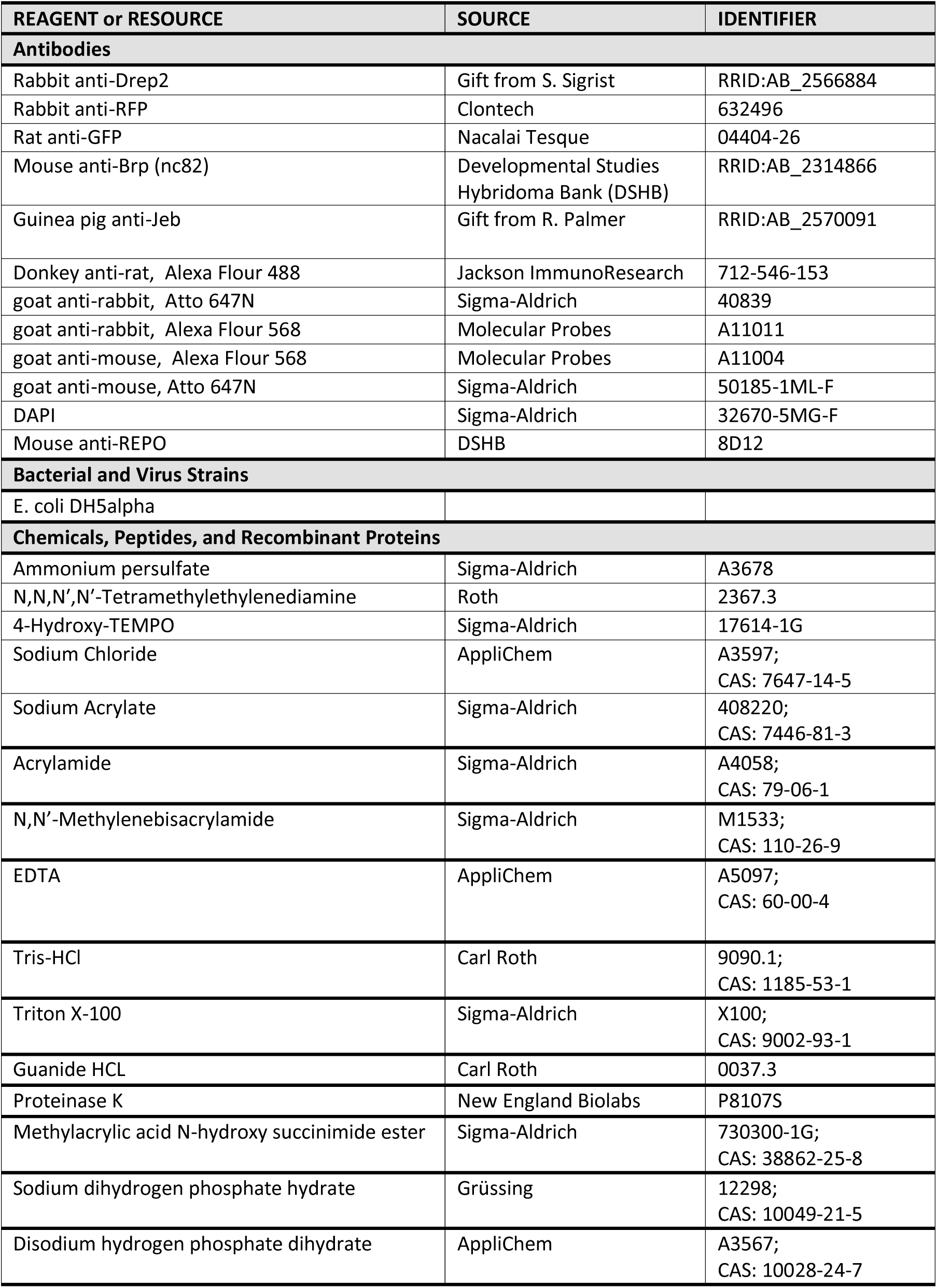
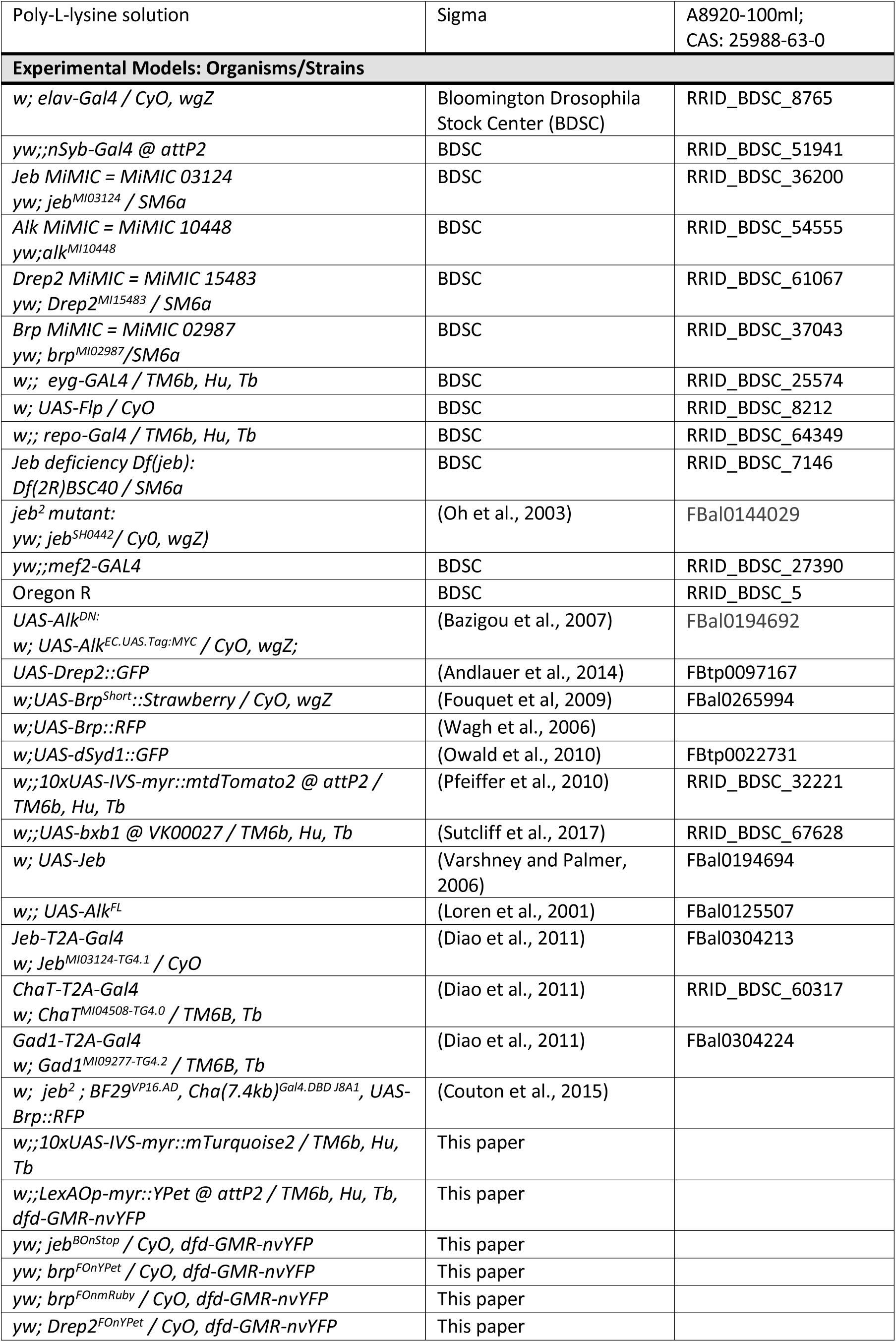
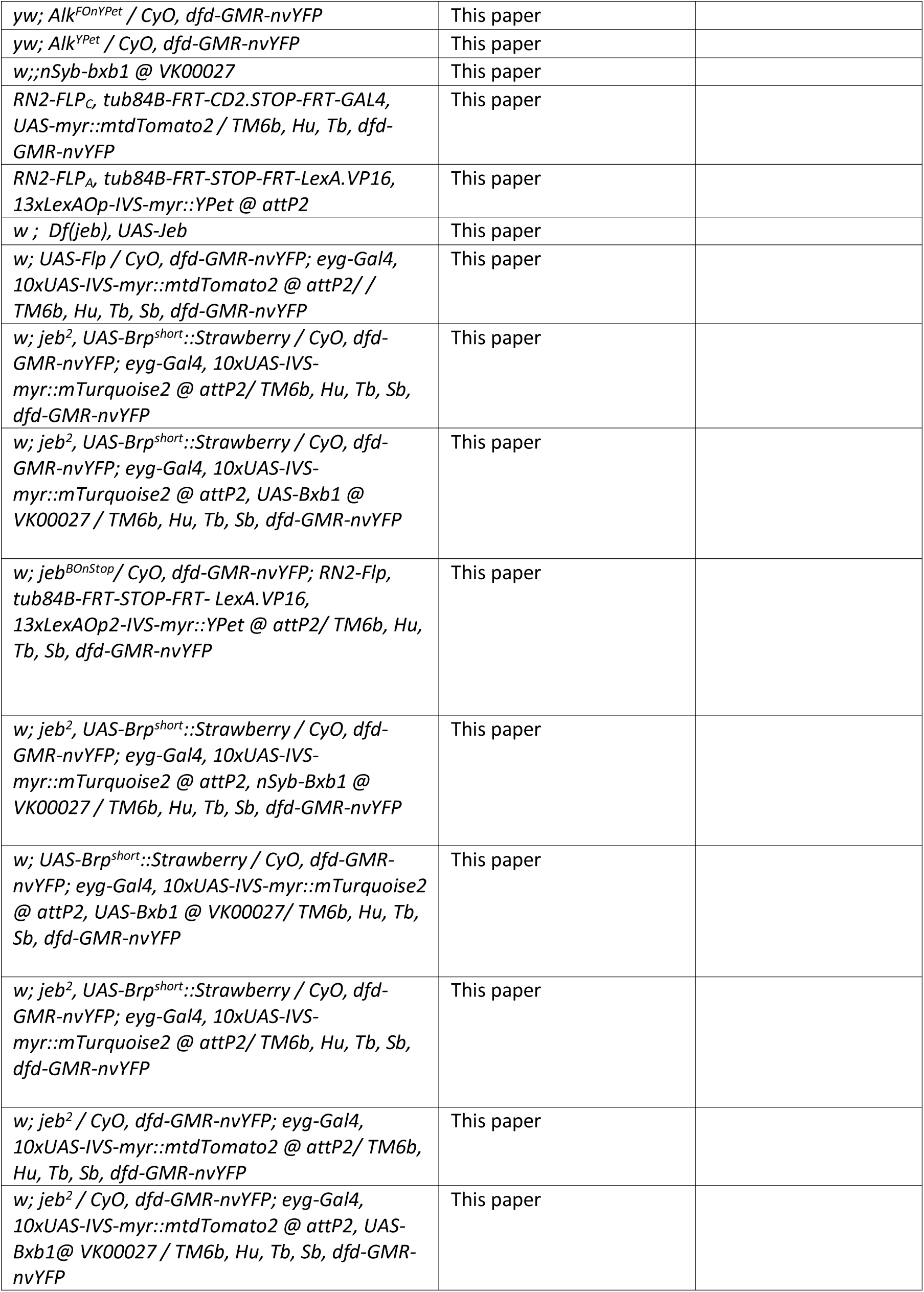
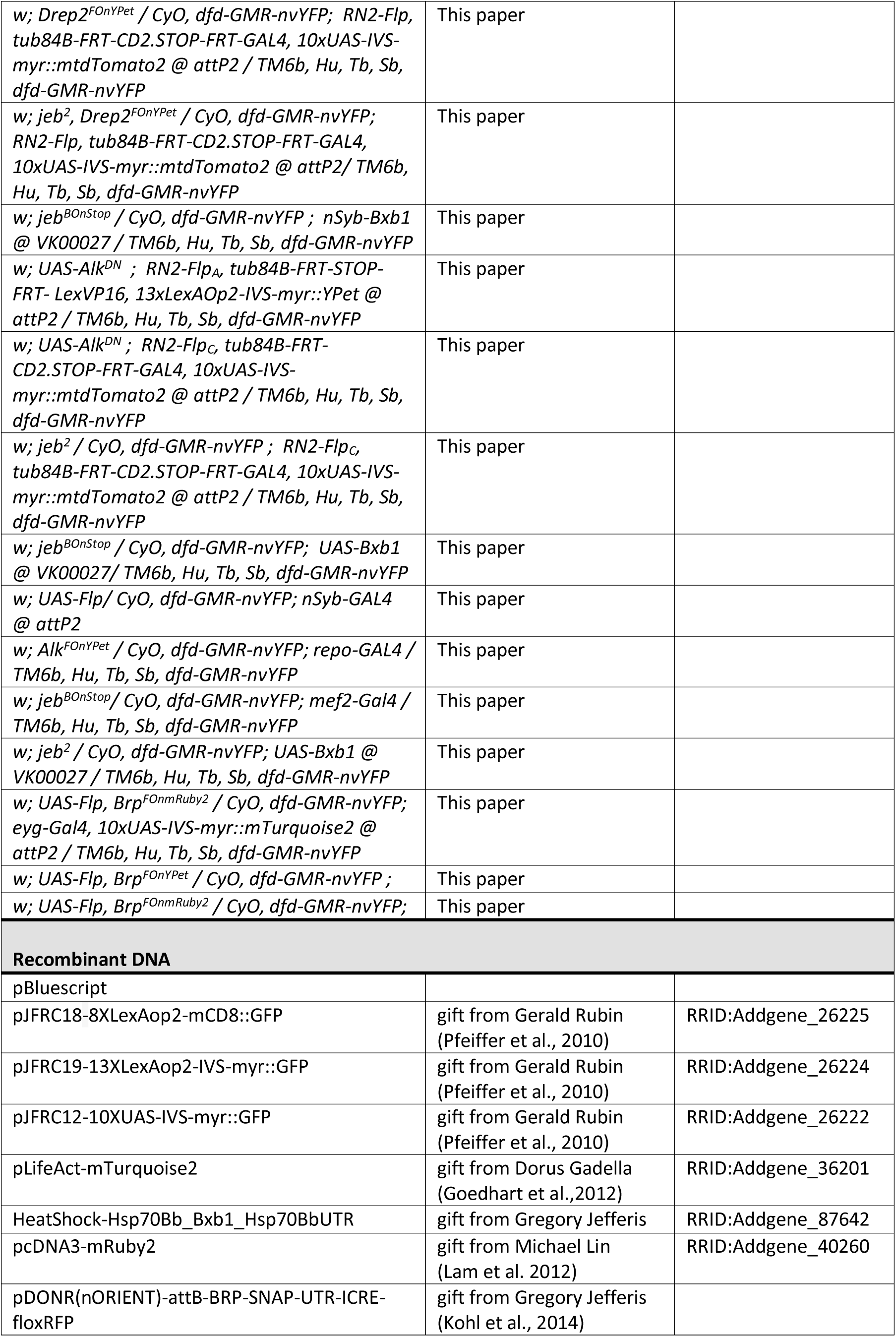
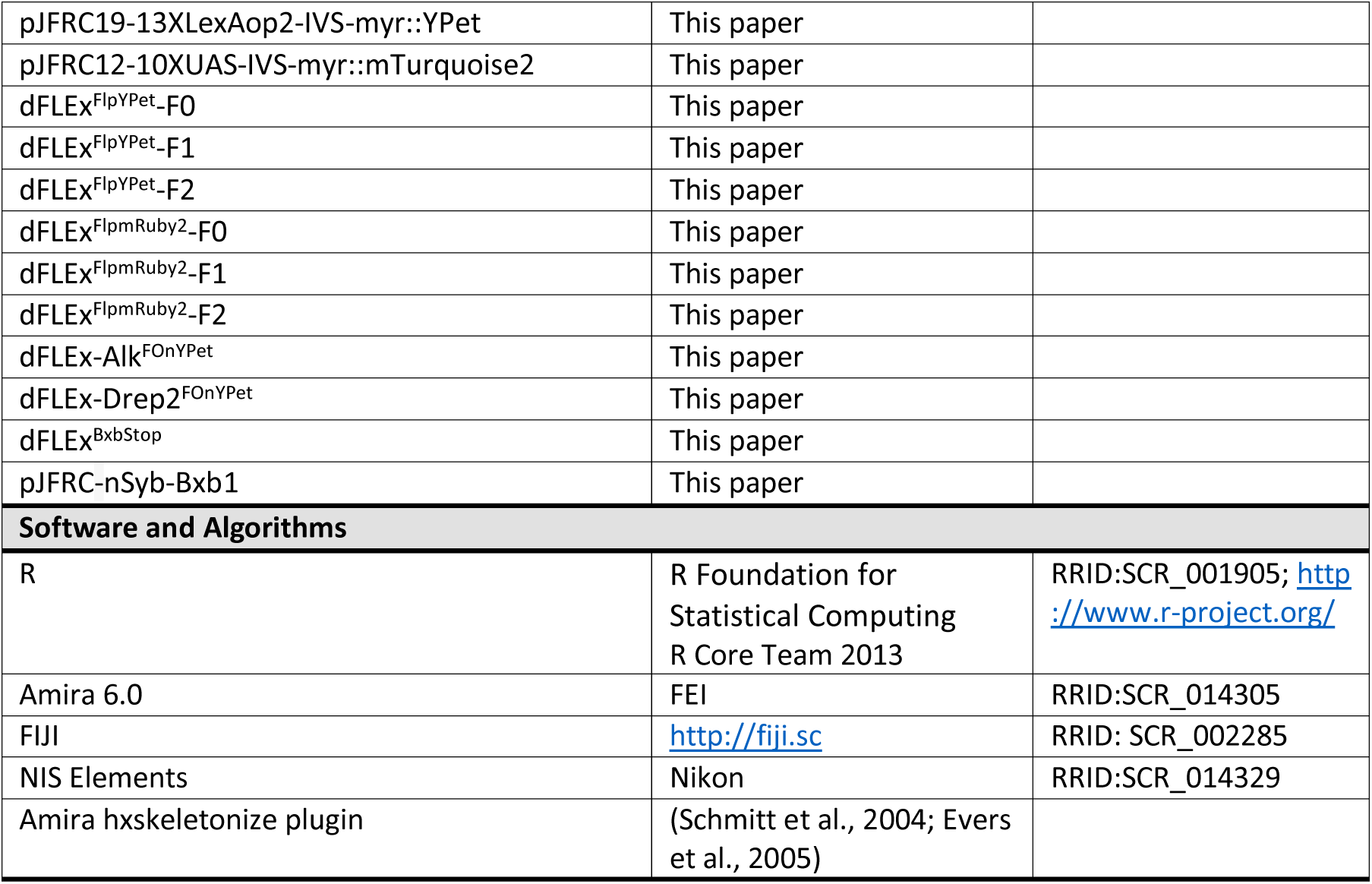
KEY RESOURCE TABLE.

### EXPERIMENTAL MODEL AND SUBJECT DETAILS

Drosophila melanogaster wild-type strain Oregon R (OrR) and mutant flies were raised on standard medium at 25°C and 60% humidity in constant darkness.

### CONTACT FOR REAGENT AND RESOURCE SHARING

Further information and requests for resources and reagents should be directed to and will be fulfilled by the Lead Contact, Jan Felix Evers

## METHOD DETAILS

### Staging of animals

Flies were allowed to lay eggs on apple juice-based agar medium overnight at 25 °C, or 30 °C for experiments using the RN2-FLP alleles (Ou et al., 2008). To aid collection of trachea-filled embryos, the chorion was removed by incubation with household bleach for 2 minutes. Larvae that hatched within the next hour were collected for data collection (designated 0 h ALH), or allowed to develop further in yeast paste until the desired age.

### Intra-vital imaging

Larvae were removed from food and rinsed with tap water before placing them onto a cover slip in a custom build imaging chamber with the ventral side facing the microscope objective. Animals were anaesthetized with 18% desflurane (Baxter) (Füger et al., 2007), vaporized with a D-Vapor apparatus (Dräger Medical). A piezo motor inside the imaging chamber allowed to gently compress the anesthetized larvae with a second glass cover slip to improve optical contact and thus image quality. Image stacks were acquired in intact, transiently anesthetized first (0h ALH), and second (24 h ALH) instar stage. At third instar (48 h ALH), the larval CNS was dissected in Sørensen buffer and acutely imaged without fixation.

### Immunohistochemistry

Embryonic and larval VNCs were dissected in Sørensen’s phosphate buffer (pH 7.2, 0.075 M). Samples were mounted on a poly-L-lysine (Sigma) coated coverslip and fixed for 15 minutes in 2% paraformaldehyde (v/v) (Electron Microscopy Sciences) and 3% sucrose (w/v) in Sørensen’s. After 30 minutes of washing in buffer containing 0.3% Triton-X-100 (Sigma-Aldrich), primary antibody treatment was performed overnight at 10°C. After a minimum of 30 minutes of washing specimen were incubated with secondary antibodies for 2 hours at room temperature. The following antibodies were used: Primary: rabbit anti-Drep2 1:500 (S. Sigrist), rabbit anti-RFP 1:500 (Clontech), anti-GFP 1:1000 (Nacalai Tesque), nc82 1:700 (DSHB), anti-Jeb 1:2000 (R. Palmer). Secondary antibodies: donkey anti-rat Alexa488 1:500 (Jackson ImmunoResearch), goat anti-rabbit Alexa568 1:1000 (Molecular Probes), goat anti-rabbit atto647N 1:500 (Sigma-Aldrich), goat anti-mouse Alexa568 1:1000 (Molecular Probes), goat anti-mouse alexa647N 1:1000 (Sigma-Aldrich).

### Expansion microscopy of ventral nerve cord preparations

To perform expansion microscopy, samples were immunostained as described above, followed by incubation in 1 mM Methacrylic acid *N*-hydroxysuccinimide ester for 1 hour at RT (Chozinski et al., 2016). To minimize tissue warping in high saline buffer, samples were incubated in 30% and 60% monomer solution (MS) (1xPBS, 2 M NaCl, 2.5% (wt/wt) acrylamide, 0.15% (wt/wt) N,N′-methylenebisacrylamide, 8.625% (wt/wt) sodium acrylate) for 15 minutes each and 100% MS for 45 minutes at 4°C. Gelling was performed at 37 °C for 1h after adding ammonium persulfate (Sigma-Aldrich), N,N,N′,N′-Tetramethylethylenediamine (Roth) and 4-Hydroxy-TEMPO (Sigma-Aldrich) to MS. After gelling, excess gel was removed and embedded specimen were placed in digestion buffer (1X TAE, 0.5% Triton X-100, 0.8 M guanide HCL) with 8 units/ml Proteinase K (NEB) for 2 hours at 37 °C. The gel was expanded in deionized water for a total of 1h. Water was exchanged every 15 min. Gels were mounted on poly-L-lysine coated cover slips for imaging.

### Image Acquisition

All images were acquired with a custom-built spinning disk confocal field scanner mounted on a Nikon Ti microscope, using a 60x/1.2 N.A. Nikon water immersion objective. Images were acquired with a Photometrics Evolve Delta camera at an effective voxel size of 0.267 × 0.267 × 0.300 μm. Image acquisition was controlled by MicroManager (NIH, Arthur et al, 2014) or NIS Elements (Nikon).

### Electrophysiology and Electroshock assay

Whole-cell patch clamp recordings from RP2 motoneurons were performed at 48 h and 0-3 h ALH and analyzed as previously described (Baines and Bate, 1998). Electroshock assay was performed as previously described (Marley and Baines, 2011). A 2.3 V DC pulse for 2 s, created by a constant voltage generator (DS2A-mkII, Digitimer Ltd., Welwyn Garden City, Hertfordshire, UK), was applied on the anterior-dorsal cuticle of third instar larvae. The time to resumption of normal crawling behavior was measured as recovery time (RT).

### Cloning

All transformation plasmid were made by Gibson Assembly (Gibson et al., 2009). The dFLEx^FlpYPet^ linear fragment (Fig S4) was made by gene synthesis; FRT and FRT_F5_ sequences from (Schlake and Bode, 1994) and TN5 linker sequence from (Matkovic et al., 2013; Sheridan et al., 2002) and YPet sequence from (Nguyen and Daugherty, 2005). Figures S4–S7 show the primers used to make each plasmid. pJFRC12-10XUAS-IVS-myr::mTurquoise2 was made my amplifying mTurquoise2 from LifeAct-mTurquoise2 with Gibson compatible overlaps for insertion into pJFRC12-10XUAS-IVS-myr::GFP, directly replacing GFP. pJFRC19-13XLexAop2-IVS-myr::YPet was generated by subcloning YPet from dFLEx^FlpYPet^-F0 into pJFRC19-13XLexAop2-IVS-myr::GFP, directly replacing GFP.

### Transgenic animals

Transgenic flies were made at University of Cambridge, Department of Genetics Fly Facility. Flies carrying *brp^FOnYPet^* and *brp^FOnmRuby2^* were generated by insertion of dFLEx^FlpYPet^-F0 and dFLEx^FlpmRuby2^-F0 respectively into Brp^MI02987^, located in a coding intro between exon 9 and 10. Drep2^FOnYPet^ flies were made by insertion of dFLEx-Drep2^FOnYPet^ into *Drep2^MI15483^* followed by induced homologous recombination by I-CreI expression in the germ line. The induced Drep2^FOnYPet^ allele labels Drep2 at the N-Term. Alk^FOnYPet^ flies resulted from insertion of dFLEx-Alk^FOnYPet^ into Alk^MI10448^ followed by I-CreI induced homologous recombination in the germ line. Induced Alk^FOnYPet^ labels Alk at the C-Term. The *jeb^BOnStop^* allele resulted from dFLEx-Jeb^BOnStop^ insertion into Jeb^MI03124^, immediately preceding the region coding for the type A LDL receptor domain. pJFRC18-nSyb-Bxb1 was inserted into VK00027 to generate *nSyb-Bxb1* flies. pJFRC19-13XLexAop2-IVS-myr::YPet and pJFRC12-10XUAS-IVS-myr::mTurquoise2 were injected into attP2.

## QUANTIFICATION AND STATISTICAL ANALYSIS

### Analysis of neuronal morphology

The branched morphology of neurons was quantified by digital 3D reconstruction using the hxskeletonize plugin (Evers et al., 2005; Schmitt et al., 2004) with Amira 6.0 (FEI). Dendritic and axonal growth dynamics were annotated manually by comparing 3D reconstructions from adjacent time points of intra-vital imaging sequences. Identification of branches was started from the primary neurite following the largest branches until finally comparing thin, terminal structures. Following criteria were evaluated: (1) position of the branch origin (branchpoint) along the parent branch; (2) position of the branch insertion relative to other branches along the parent branch; (3) insertion and growth direction of branch relative to the parent branch, primary neurite, and cell body; (4) shape of the branch (curvature/bend); (5) length of the branch.

The recording angle differed between subsequent intra-vital imaging time points of the same neuron. Limited by the optical resolution of spinning disk confocal microscopy, we found that insertion angles of obviously identical branches into their parent branches vary by up to 90 degrees between time points. Therefore, changes of insertion angle up to 90 degrees were considered a modification of an existing branch; variations larger than 90 degrees were indicative for a newly formed branch. We found neuronal branches that were embryonically established and maintained until 48 h ALH to scale by a factor of 1.3 from 0-24 h ALH, and 1.4 from 24-48 h ALH. We applied the according scale factor to help distinguish scaling growth from tip growth: sections of individual branches that grew longer than what could be assumed by scaling growth were classified as new length. Data was exported as csv and processed using customized R scripts.

### Quantification of synaptic contacts

Image stacks of RP2 motoneurons were recorded with expansion microscopy, and processed in FIJI by applying a Gaussian 3D filter, and subtracting background noise using the rolling ball algorithm. Drep2 puncta were manually counted as mature synapses if found to juxtapose Brp (anti-Brp), using the FIJI Cell Counter plugin (http://rsbweb.nih.gov/ij/plugins/cell-counter.html). Profiles of fluorescence intensity across synaptic contact were determined along a straight line connecting Drep2 and Brp signals, and averaged between multiple samples. Distances were measured according to microscope magnification, and corrected by the experimentally determined expansion factor of 3.7.

### Statistical Analysis

All statistical comparisons were made using pairwise Student’s t test for multiple comparison of parametric data followed by p-adjustment after Holm, unless otherwise noted. Statistical tests were performed using R (R-project). Data are reported as mean ± standard error of mean (SEM). Dots represent single data points. Plots were generated with ggplot2 (NB: https://cran.r-project.org/web/packages/ggplot2/citation.html)

## DATA AND SOFTWARE AVAILABILITY

All data published in this study are available upon request.

---

**Figure S1.**
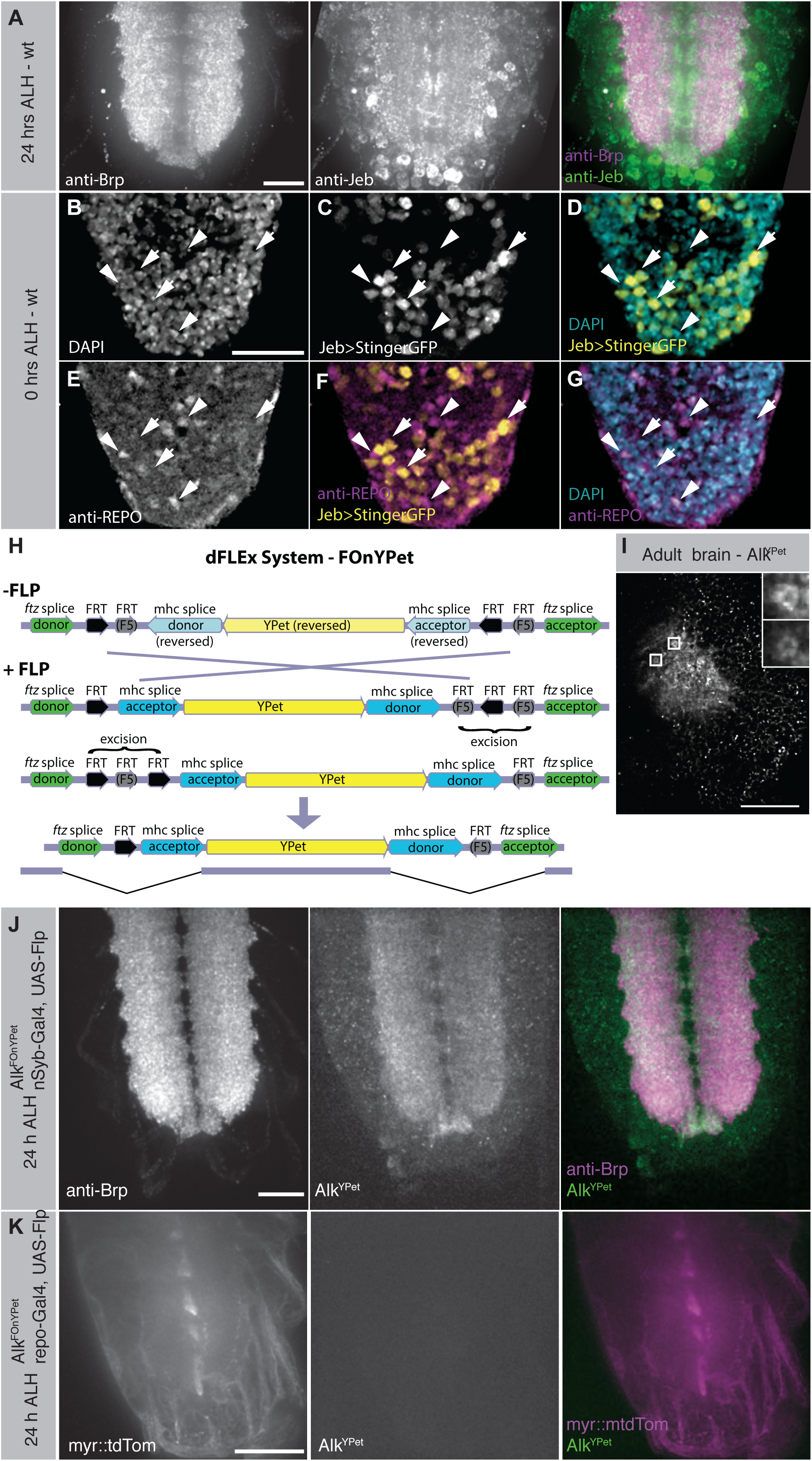
Jeb and Alk are expressed in neurons, but not glia. A – Jeb (anti-Jeb) strongly localizes to the neuropile (α -Brp), and more weakly to cell bodies. B-G –Jeb is expressed in neurons, but not glia. *jeb* translational activity (*Jeb-T2A-Gal4*) is reported by GFP (yellow; *UAS-StingerGFP*) localizing to nuclei (DAPI, cyan). Jeb translation is evident in neurons (arrows), but not glial cells (anti-REPO, arrowheads). H – Schematic of the dFLEx system depicting the FOnYPet construct (FLPase dependent on-switchable YPet fluorophore). A YPet flourophore is flanked by splice acceptor and donor as well as orthogonal FRT and FRT5 sites. FLPase expression induces DNA inversion between corresonding FRT or FRT5 sites; this inversion is subsequently stabilized by DNA excision between corresponding FRT sites. I – Constitutive, endogenously labeled Alk^YPet^ (after FLP event in germline) reveals Alk localization to postsynaptic structures of microglomeruli (insets) in the mushroom body calyx. J – Induction of Alk^FOnYPet^ in all neurons reveals strong Alk^Ypet^ (green) localization to the neuropile (anti-Brp, magenta), and weaklier to neuronal cell bodies. K – Induction of Alk^FOnYPet^ in glia cells (repo>myr::mtdTom) does not yield detectable levels of Alk^YPet^ in the ventral nerve cord. Scale bar = 20 µm (A, B-G, J, K), 100 µm (I)

**Figure S2.**
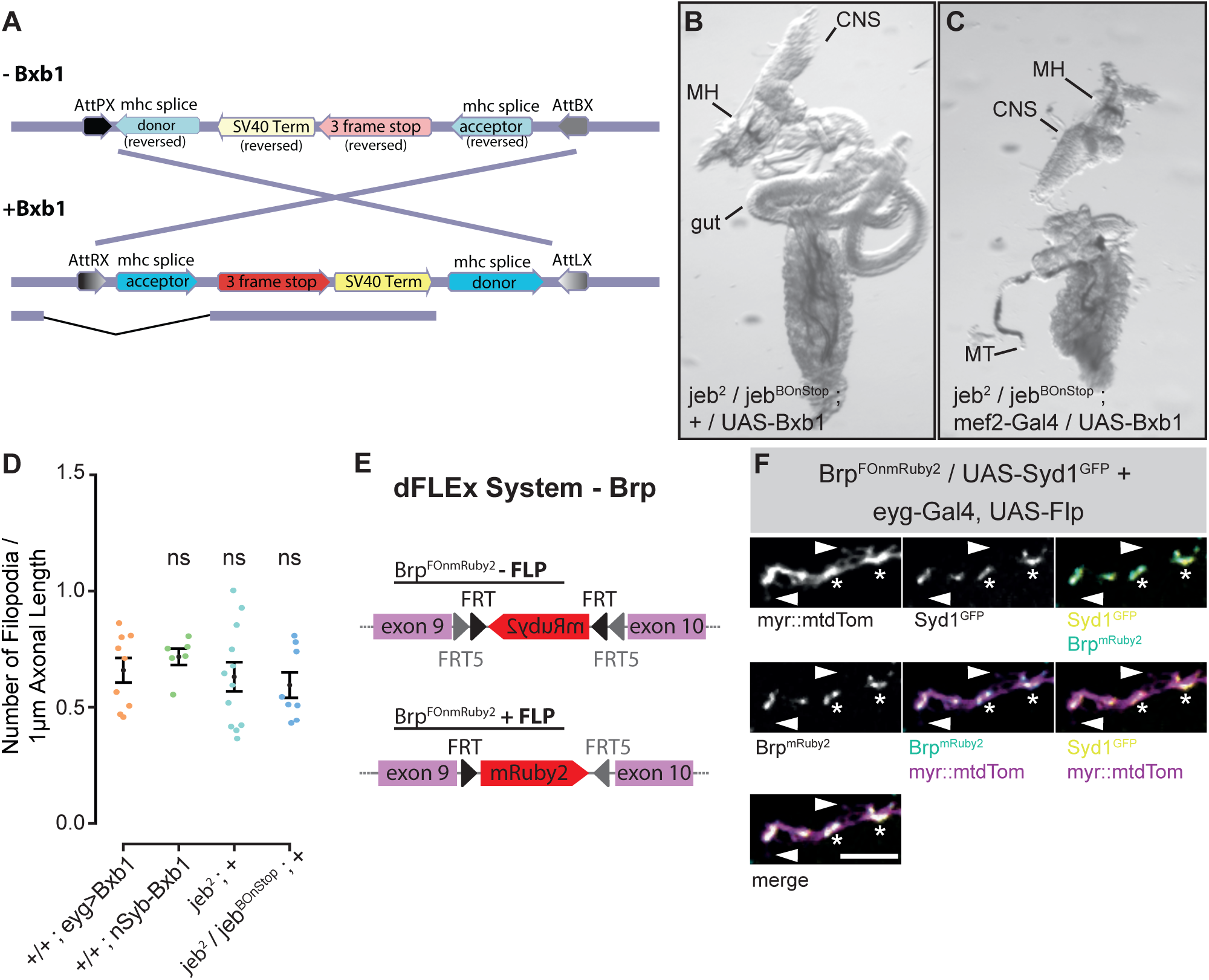
A – The BOnStop construct consist of a Stop codon in all 3 frames and an SV40 termination sequence, flanked by mhc splice acceptor/donor and AttPX / AttBX sites. Bxb1 -integrase cuts at AttPX /AttBX sites and locks the cassette in inverted orientation by conversion of AttPX / AttBX to AttRX / AttLX sequences. When placed in a gene’s coding intron, Bxb1 expression thus induces targeted mutation by premature termination of transcription and translation. B,C – (B) Transheterozygous combination of the *jeb^BOnStop^* allele over *jeb^2^* allows normal gut development. (C) Induction of *jeb^BOnStop^* to the mutant *jeb^Stop^* allele by Bxb1 expression in the early mesoderm (*mef2>bxb1*) reliably disrupts gut formation. D – Quantification of the numbers of filopodia along IN_lat_ axons confirming that increased filopodial growth in induced *jeb^BOnStop^* condition is specific to loss of Jeb (Fig 2B-G), and not caused by individual components of the BOnStop technique. E - Schematic of the Brp^FOnmRuby^ allele. The FOnmRuby construct is inserted between *brp* exons 9 and 10 (magenta) at the MiMIC site MI02987. F – Co-expression (eyg-Gal4, UAS-Flp) of Brp^FOnmRuby2^ (cyan) and UAS-Syd1^GFP^ (yellow) in IN_lat_ (magenta). Syd1 and Brp colocalize within the axon (asteriks) but do not locate to filopodia (arrowhead), demonstrating that neither immature nor mature presynaptic release sites form in presynaptic filopodia.

**Figure S3.**
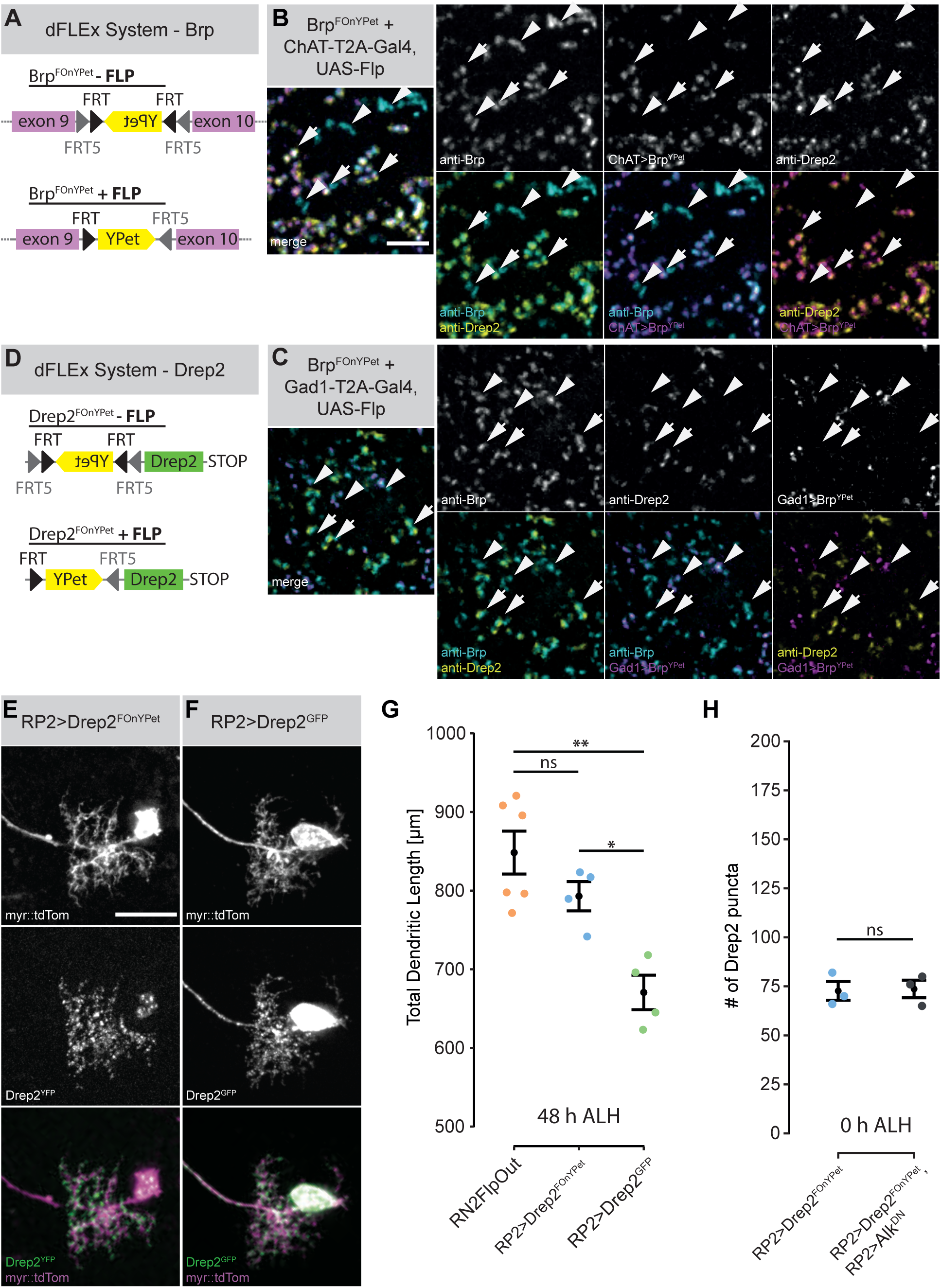
Endogenousy labeled Drep2^FOnYPet^ does not affect dendritic growth. A - Schematic of the Brp^FOnYpet^ allele. B,C – Expansion microscopy demonstrates that (B) cholinergic presynaptic specializations (arrows; Brp^FOnYPet^,ChAT>UAS-Flp) appose Drep2 profiles (anti-Drep2, yellow). Other types of presynaptic specializations (arrowheads, anti-Brp, cyan) are not paired with Drep2 profiles. (C) Brp^YPet^ expressed in GABAergic neurons (Brp^FOnYPet^,Gad1> Flp) does not appose Drep2 puncta (anti-Drep2, yellow). D - Schematic of the Drep2^FOnYpet^ allele. FLP expression induces endogenous Drep2 labeling with a YPet flourophore (yellow) at the 5’-end of Drep2 (green). E,F – Comparison of RP2 motorneuron dendrites (myr::mtdTom), (E) with endogenously labeled Drep2^YPet^ (Drep2^FOnYPet^,RP2>FLP), and (F) overexpression of Drep2^GFP^. Overexpression of Drep2^GFP^ reports artefactually high concentration of Drep2 in the cell body, primary neurite, axon and throughout the dendritic arbor compared to endogenous Drep2^YPet^. G – Quantification of RP2 dendritic length in control, endogenously labeled Drep2^YPet^ (RP2>Drep^FOnYPet^) and Drep2^GFP^ overexpression (RP2>Drep2^GFP^) reveals that overexpression of Drep2^GFP^ causes dendritic undergrowth that can be circumvented using endogenously tagged Drep2^YPet^. H – Quantification of mature postsynaptic specialization (Drep2^YPet^ apposed to anti-Brp profiles) at 0 h ALH shows that embryonic development of postsynaptic specialization is not regulated by Alk signalling. Welch two-sample t-test. Scale bars = 10 µm (B,C), 20 µm (E). *p<0.05,**p<0.01, ns – not significant

**Figure S4:**
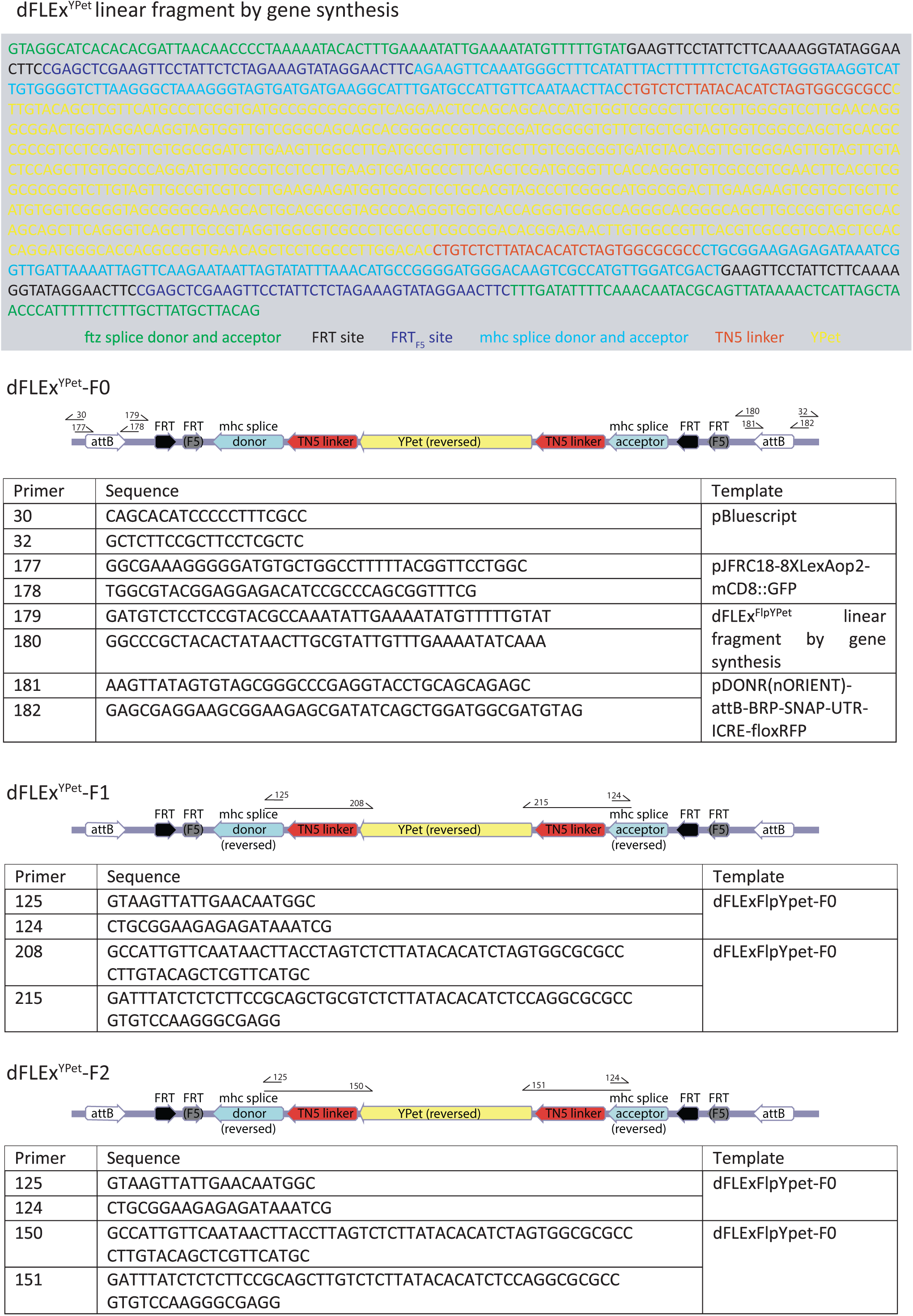
dFLEx^FlpYPet^ related to STAR methods

**Figure S5:**
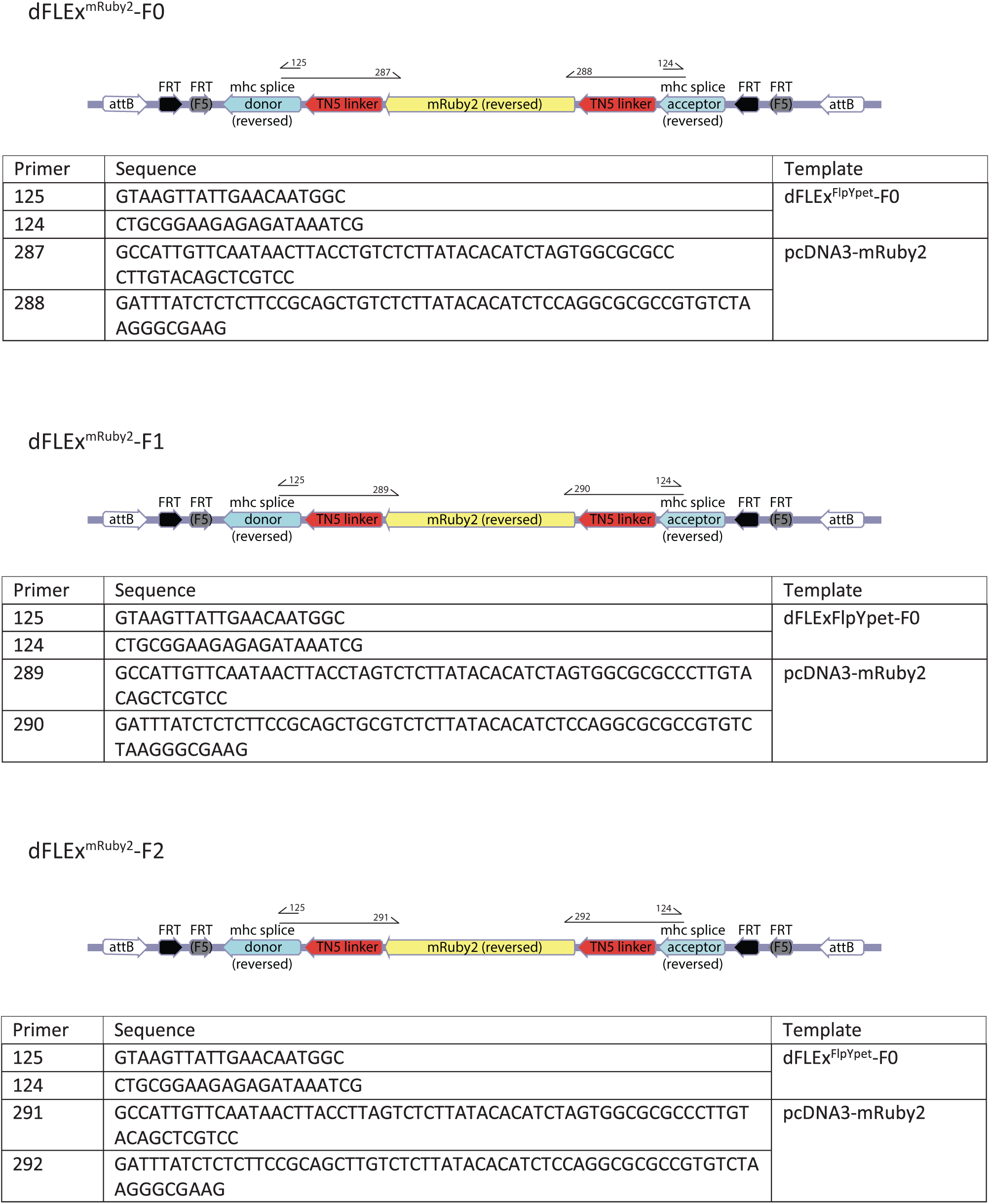
dFLEx^FlpmRuby2^ related to STAR methods

**Figure S6:**
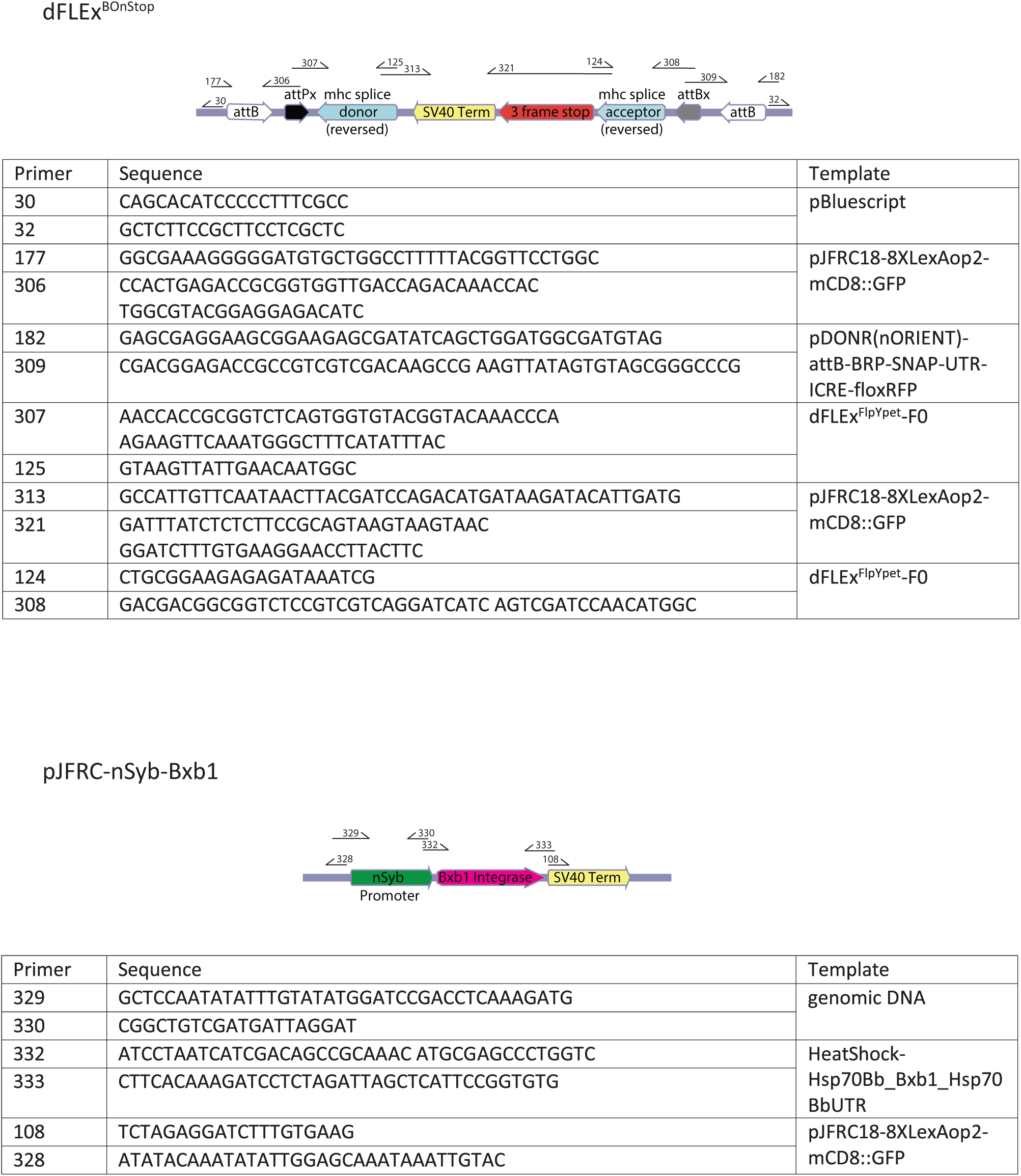
dFLEx^BOnStop^ and pJFRC-nSyb-Bxb1 related to STAR methods

**Figure S7:**
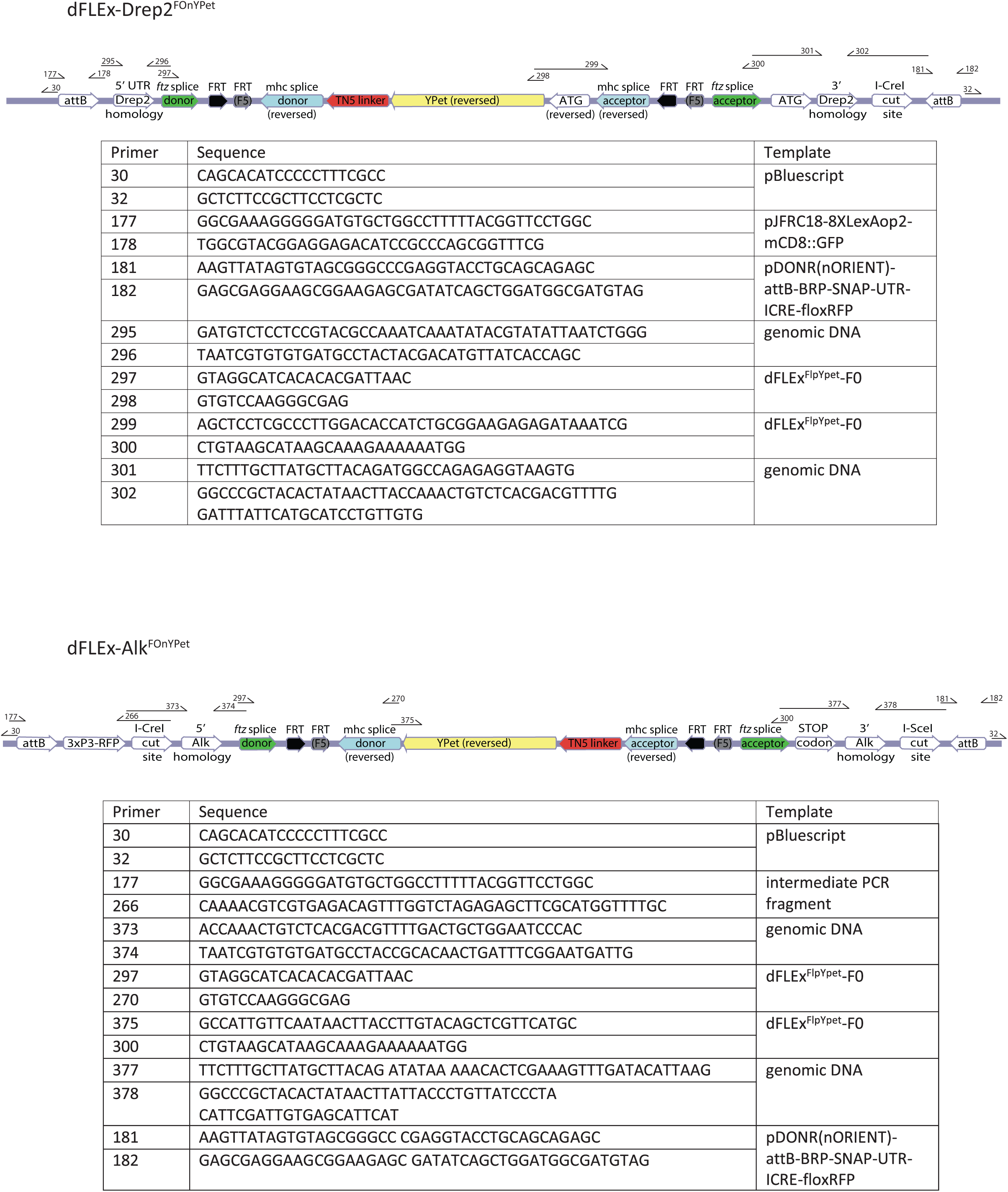
dFLEx-Drep2^FOnYPet^ and dFLEx-Alk^FOnYPet^ related to STAR methods

**Table S1:** Experimental genotypes

| Figure | Genotype |
| --- | --- |
| **Figure 1** |  |
| 1B | *w; jeb<sup>2</sup> / Df(jeb), UAS-Jeb ; BF29<sup>VP16.AD</sup>, Cha(7.4kb)<sup>Gal4.DBD</sup> J8A1, UAS-Brp::RFP / +* |
| 1D-E | *w; Alk<sup>FOnYPet</sup> / UAS-Flp ; eyg-Gal4, 10xUAS-IVS-myr::mtdTomato2 @ attP2 / +* |
| 1F | *w; Alk<sup>FOnYPet</sup> / + ; RN2-Flp, tub84B-FRT-CD2.STOP-FRT-Gal4, 10xUAS-IVS-myr::mtdTomato2 @ attP2 / +* |
| **Figure 2** |  |
| 2B+F-H | *w; jeb<sup>2</sup>, UAS-Brp<sup>short</sup>::Strawberry / jeb<sup>BOnStop</sup> ; eyg-Gal4, 10xUAS-IVS-myr::mTurquoise2 @ attP2 / RN2-Flp, tub84B-FRT-STOP-FRT- LexA.VP16, 13xLexAOp2-IVS-myr::YPet @ attP2* |
| 2C+F-H | *w; jeb<sup>2</sup>, UAS-Brp<sup>short</sup>::Strawberry / jeb<sup>BOnStop</sup>; eyg-Gal4, 10xUAS-IVS-myr::mTurquoise2 @ attP2, UAS-Bxb1 @ VK00027 / RN2-Flp, tub84B-FRT-STOP-FRT- LexA.VP16, 13xLexAOp2-IVS-myr::YPet @ attP2* |
| 2D+F-H | *w; jeb<sup>2</sup>, Brp<sup>short</sup>::Strawberry / jeb<sup>BOnStop</sup> ; eyg-Gal4, 10xUAS-IVS-myr::mTurquoise2 @ attP2, nSyb-Bxb1 @ VK00027 / RN2-Flp, tub84B-FRT-STOP-FRT- LexA.VP16, 13xLexAOp2-IVS-myr::YPet @ attP2* |
| 2E+F-G | *w; UAS-Brp<sup>short</sup>::Strawberry / UAS-Alk<sup>DN</sup> ; eyg-Gal4, 10xUAS-IVS-myr::mTurquoise2 @ attP2, UAS-Bxb1 @ VK00027 / RN2-Flp<sub>A</sub>, tub84B-FRT-STOP-FRT- LexA.VP16, 13xLexAOp2-IVS-myr::YPet @ attP2* |
| **Figure 3** |  |
| 3A-D | *w; jeb<sup>2</sup> / jeb<sup>BOnStop</sup> ; eyg-Gal4, 10xUAS-IVS-myr::mtdTomato2 @ attP2 / +* |
| 3C+D | *w; jeb<sup>2</sup> / jeb<sup>BOnStop</sup> ; eyg-Gal4, 10xUAS-IVS-myr::mtdTomato2 @ attP2, UAS-Bxb1 @ VK00027 / +* |
| **Figure 4** |  |
| 4A-E+H | *w; Drep2<sup>FOnYPet</sup> / + ; RN2-Flp, tub84B-FRT-CD2.STOP-FRT-GAL4, 10xUAS-IVS-myr::mtdTomato2 @ attP2 / +* |
| 4F+H | *w; Drep2<sup>FOnYPet</sup> / UAS-Alk<sup>DN</sup> ; RN2-Flp, tub84B-FRT-CD2.STOP-FRT-GAL4, 10xUAS-IVS-myr::mtdTomato2 @ attP2 / +* |
| 4G+H | *w; jeb<sup>2</sup>, Drep2<sup>FOnYPet</sup> / jeb<sup>BOnStop</sup> ; RN2-Flp, tub84B-FRT-CD2.STOP-FRT-GAL4, 10xUAS-IVS-myr::mtdTomato2 @ attP2 / nSyb-Bxb1 @ VK00027* |
| 4H | *w; jeb<sup>2</sup>, Drep2<sup>FOnYPet</sup> / + ; RN2-Flp, tub84B-FRT-CD2.STOP-FRT-GAL4, 10xUAS-IVS-myr::mtdTomato2 @ attP2 / nSyb-Bxb1 @ VK00027* |
| **Figure 5** |  |
| 5A,C-G,I-K | *w; jeb<sup>2</sup> / jeb<sup>BOnStop</sup> ; RN2-Flp, tub84B-FRT-CD2.STOP-FRT-GAL4, 10xUAS-IVS-myr::mtdTomato2 @ attP2 / +* |
| 5B-F,H-K | *w; jeb<sup>2</sup> / jeb<sup>BOnStop</sup> ; RN2-Flp, tub84B-FRT-CD2.STOP-FRT-GAL4, 10xUAS-IVS-myr::mtdTomato2 @ attP2 / nSyb-Bxb1 @ VK00027* |
| **Figure 6** |  |
| 6A+E | *w; + / + ; RN2-Flp<sub>A</sub>, tub84B-FRT-STOP-FRT- LexVP16, 13xLexAOp2-IVS-myr::YPet @ attP2 / +* |
| 6B+E | *w; jeb<sup>2</sup> / +; RN2-Flp<sub>A</sub>, tub84B-FRT-STOP-FRT- LexVP16, 13xLexAOp2-IVS-myr::YPet @ attP2 / nSyb-Bxb1 @ VK00027* |
| 6C+E | *w; jeb<sup>2</sup> / jeb<sup>BOnStop</sup> ; RN2-Flp<sub>A</sub>, tub84B-FRT-STOP-FRT- LexVP16 13xLexAOp2-IVS-myr::YPet @ attP2 / nSyb-Bxb1 @ VK00027* |
| 6D+E | *w; elav-Gal4 / UAS-Alk<sup>DN</sup> ; RN2-Flp<sub>A</sub>, tub84B-FRT-STOP-FRT- LexVP16, 13xLexAOp2-IVS-myr::YPet @ attP2 / +* |
| 6F+J | *w; +/+; RN2-Flp<sub>C</sub>, tub84B-FRT-CD2.STOP-FRT-GAL4, 10xUAS-IVS-myr::mtdTomato2 @ attP2 / +* |
| 6G+J | *w; +/UAS-Alk<sup>DN</sup>; RN2-Flp<sub>C</sub>, tub84B-FRT-CD2.STOP-FRT-GAL4, 10xUAS-IVS-myr::mtdTomato2 @ attP2 / +* |
| 6H+J | *w; +/UAS-Alk<sup>DN</sup>; RN2-Flp<sub>C</sub>, tub84B-FRT-CD2.STOP-FRT-GAL4, 10xUAS-IVS-myr::mtdTomato2 @ attP2 / UAS-Alk<sup>FL</sup>* |
| 6I+J | *w; jeb<sup>2</sup> / jeb<sup>BOnStop</sup>; RN2-Flp<sub>C</sub>, tub84B-FRT-CD2.STOP-FRT-GAL4, 10xUAS-IVS-myr::mtdTomato2 @ attP2 / UAS-Bxb1 @ VK00027* |
| **Figure 7** |  |
| 7A-G | *w; +/+; RN2-Flp<sub>C</sub>, tub84B-FRT-CD2.STOP-FRT-GAL4, 10xUAS-IVS-myr::mtdTomato2 @ attP2 / +* |
| 7C-G | *w; +/UAS-Alk<sup>DN</sup>; RN2-Flp<sub>C</sub>, tub84-FRT-STOP-FRT-Gal4, 10xUAS-IVS-myr::mtdTomato2 @ attP2 / +* |
| 7C-G | *w; jeb<sup>2</sup> / jeb<sup>BOnStop</sup>; RN2-Flp<sub>C</sub>, tub84-FRT-STOP-FRT-Gal4, 10xUAS-IVS-myr::mtdTomato2 @ attP2 / nSyb-Bxb1 @ VK00027* |
| **Figure S1** |  |
| S1A | *Oregon R* |
| S1B-G | *w; jeb-T2A-Gal4 / UAS-StingerGFP; +/+* |
| S1I | *w, Alk<sup>YPet</sup> / +; +/+* |
| S1J | *w; Alk<sup>FOnYPet</sup> / UAS-Flp; nSyb-GAL4 @ attP2* |
| S1K | *w; Alk<sup>FOnYPet</sup> / UAS-Flp; repo-GAL4* |
| **Figure S2** |  |
| S2B | *w; jeb<sup>2</sup> / jeb<sup>BOnStop</sup>; UAS-Bxb1 @ VK00027 / +* |
| S2C | *w; jeb<sup>2</sup> / jeb<sup>BOnStop</sup>; mef2-Gal4 / UAS-Bxb1 @ VK00027* |
| SD+E | *w; UAS-Brp<sup>short</sup>::Strawberry / +; eyg-Gal4, 10xUAS-IVS-myr::mTurquoise2 @ attP2, UAS-Bxb1 @ VK00027 / +*<br>*w; UAS-Brp<sup>short</sup>::Strawberry / +; eyg-Gal4, 10xUAS-IVS-myr::mTurquoise2 @ attP2, nSyb-Bxb1 @ VK00027 / +*<br>*w; jeb<sup>2</sup>, UAS-Brp<sup>short</sup>::Strawberry / +; eyg-Gal4, 10xUAS-IVS-myr::mTurquoise2 @ attP2 / +*<br>*w; jeb<sup>2</sup>, UAS-Brp<sup>short</sup>::Strawberry / jeb<sup>BOnStop</sup>; eyg-Gal4, 10xUAS-IVS-myr::mTurquoise2 @ attP2 / +* |
| S2F | *w; UAS-Flp, Brp<sup>FOnmRuby2</sup> / UAS-Syd1<sup>GFP</sup>; eyg-Gal4, 10xUAS-IVS-myr::mTurquoise2 @ attP2 / +* |
| **Figure S3** |  |
| S3B | *w; UAS-Flp, Brp<sup>FOnYPet</sup> / +; ChAT-T2A-Gal4 / +* |
| S3C | *w; UAS-Flp, Brp<sup>FOnYPet</sup> / +; Gad1-T2A-Gal4 / +* |
| S3E+G | *w; Drep2<sup>FOnYPet</sup> / +; RN2-Flp<sub>C</sub>, tub84B-FRT-CD2.STOP-FRT-GAL4, 10xUAS-IVS-myr::mtdTomato2 @ attP2 / +* |
| S3F+G | *w; UAS-Drep2<sup>GFP</sup> / +; RN2-Flp<sub>C</sub>, tub84B-FRT-CD2.STOP-FRT-GAL4, 10xUAS-IVS-myr::mtdTomato2 @ attP2 / +* |
| S3G | *w; +/+; RN2-Flp<sub>C</sub>, tub84B-FRT-CD2.STOP-FRT-GAL4, 10xUAS-IVS-myr::mtdTomato2 @ attP2 / +* |
| S3H | *w; Drep2<sup>FOnYPet</sup> / +; RN2-Flp<sub>C</sub>, tub84B-FRT-CD2.STOP-FRT-GAL4, 10xUAS-IVS-myr::mtdTomato2 @ attP2 / +*<br>*w; Drep2<sup>FOnYPet</sup> / UAS-Alk<sup>DN</sup>; RN2-Flp<sub>C</sub>, tub84B-FRT-CD2.STOP-FRT-GAL4, 10xUAS-IVS-myr::mtdTomato2 @ attP2 / +* |

